# CD47 promotes MAPK and epithelial-to-mesenchymal transition molecular programs to drive pro-metastatic phenotypes in non-small cell lung cancer

**DOI:** 10.1101/2025.09.25.678579

**Authors:** Asa P.Y. Lau, Lilian G. Zhai, Ryunosuke Hoshi, Zackary Rousseau, Abraam Zakhary, Yin Fang Wu, Rola Saleeb, Heyu Ni, Kelsie L. Thu

## Abstract

CD47 is best known for its role in tumor immune evasion; however, studies in diverse cell models indicate that it also has cell-autonomous, tumor-promoting functions which are cell type- and context-specific. Motivated by the prognostic and therapeutic significance of CD47 and the limited knowledge regarding its roles beyond immune evasion in non-small cell lung cancer (NSCLC), we sought to define the cellular and molecular processes driven by intrinsic CD47 signaling in NSCLC. Transcriptome profiling of CD47 wildtype and knockout NSCLC cells implicated its regulation of genes enriched for signatures of MAPK signaling and epithelial-to-mesenchymal transition (EMT). A significant positive association between CD47 and MAPK/EMT expression signatures was also evident in large cohorts of NSCLC cell lines and tumor tissues. Functional studies indicated that CD47 does not regulate cell proliferation in NSCLC cells like it does in other cancer types. Instead, CD47 regulates cell adhesion and migration through an ERK and EMT axis, validating our transcriptomic findings. Moreover, CD47 loss-of-function significantly diminished the ability of NSCLC cells to metastasize *in vivo*, demonstrating the physiological relevance of cell-intrinsic CD47 signaling in lung cancer cells. Our data reveal a novel role for CD47 in relaying signals through ERK to promote EMT expression programs and pro-metastatic phenotypes in NSCLC. Although additional mechanistic studies are needed to further decipher the CD47-ERK-EMT signaling pathway, our findings reinforce the therapeutic potential of CD47, rationalizing further research to develop CD47 blockade as a multimodal therapy for NSCLC.

## 1.0 Introduction

Despite global declines in tobacco smoking rates and the implementation of screening programs in developed countries, lung cancer remains the leading cause of cancer-related deaths worldwide [1,2]. Approximately 40% of lung cancer patients are diagnosed with advanced stage disease characterized by tumor metastasis to distant sites [3]. Accordingly, outcomes for lung cancer patients are relatively poor compared to those for other common malignancies, with the 5-year overall survival rate estimated to be 27% [4]. Non-small cell lung cancer (NSCLC) is the most prevalent type of lung cancer. Current treatments include surgery, radiation, chemotherapy, targeted therapy, and/or immunotherapy which are assigned based on tumor stage, histotype, mutation status, and other biomarkers like PD-L1 [5,6]. Over the past decade, immunotherapies that inhibit the immune checkpoint proteins PD-(L)1 and CTLA-4 have shown remarkable efficacy in some NSCLC patients, however, only 30-40% of patients are eligible for these drugs and only 15-20% respond [7,8]. In NSCLC patients who respond to PD-(L)1 blockade, more than 60% develop acquired resistance [9,10]. These statistics indicate that alternative immunotherapies for treating patients ineligible for, or resistant to, current immune checkpoint inhibitors are needed to improve survival rates.

Cluster of differentiation 47, CD47 (also known as integrin associated protein, IAP), is a transmembrane protein often overexpressed on tumor cells that has emerged as a promising immunotherapeutic target in NSCLC and other cancers [11–14]. It is best known for its immunosuppressive function mediated through interactions with its receptor SIRPα, which is expressed on phagocytes like macrophages and dendritic cells. Ligation of CD47 and SIRPα stimulates a signaling cascade that transduces a potent “don’t eat me” signal to inhibit the ability of these cells to perform phagocytosis [14]. As such, cancer cells exploit this checkpoint mechanism to escape phagocytic clearance, suppress anti-tumor immune responses, and evade destruction by the immune system. Discovery of its role in immune evasion led to the development of CD47-targeted immunotherapies. Studies in preclinical models have shown that inhibiting the CD47-SIRPα checkpoint using therapeutic antibodies or fusion proteins (i.e., CD47 blockade) impairs the growth of several tumor types and that these effects are driven by relieving CD47-mediated suppression of phagocytes and their activation of adaptive tumor immunity [15].

Given its promise as an immunotherapeutic target, much research on CD47 in the oncology field has focused on its immunosuppressive function. However, apart from its role in tumor immune escape, CD47 regulates other processes that promote tumor growth and progression [14,16]. These include regulation of autophagy, cell death and survival, angiogenesis, metabolism, and response to stress, which are driven by its interactions with proteins like thrombospondin-1 (TSP-1) and integrins including αvβ3, αIIbβ3, and multiple β1 integrins [14,16]. Discoveries of these CD47 functions have been made in diverse physiological and cellular contexts including non-malignant epithelial, endothelial, and smooth muscle cells, as well as cancer models. Findings to date indicate that cell-intrinsic CD47 signaling pathways and their biological effects are pleiotropic and cell type-specific due to differential expression of CD47-interacting proteins in different cells and environments [16]. Thus, studies to decipher CD47 signaling and its cell autonomous effects specifically in lung cancer are necessary to better understand its contributions to lung tumor biology and therapeutic significance.

Investigations of CD47 in NSCLC have determined that it is frequently overexpressed and that high expression is associated with advanced tumor stage, metastasis, and poor survival [11–13,17]. Functional studies in preclinical models have shown that CD47 inhibition impairs lung tumor growth and progression [18–20], with recent work indicating that combinations of CD47 blockade with EGFR/KRAS/ALK-targeted therapies may be effective treatment strategies for NSCLC [21,22]. Associations between CD47 and cell migration, metastasis and resistance to EGFR-targeted therapy have also been reported in lung cancer [13,20,21,23,24], but the signaling mechanisms underlying CD47’s role in these processes are poorly defined.

In the present study, we sought to investigate cell-intrinsic CD47 signaling mechanisms and the tumor-promoting phenotypes they influence in NSCLC cells. Using transcriptomic profiling and functional studies in NSCLC cells engineered with and without CD47 loss of function, we discovered that CD47 promotes molecular programs that drive epithelial-to-mesenchymal transition (EMT) and pro-metastatic phenotypes, likely through an ERK-dependent signaling mechanism. Our findings provide new insights into the cell-intrinsic functions of CD47 in NSCLC, and suggest that its therapeutic potential extends beyond its immunosuppressive activity. As such, continued efforts to develop CD47 blockade as a multimodal treatment strategy for NSCLC are warranted.

## 2.0 Methods

### 2.1 Cell culture and lentivirus production

Lewis Lung Carcinoma cells stably expressing luciferase (LLC; CVCL_4358) were purchased from Promega. CMT167 (CMT) cells were purchased from Sigma (#10032302). NCI-H1299 (H1299; CVCL_0060) and NCI-H2122 (H2122; CVCL_1531) were generous gifts from the late Dr. Adi Gazdar. A549 (CVCL_0023) was obtained from the The American Type Culture Collection (#CCL-185). Cell lines were grown at 37°C and 5% CO_2_ in DMEM (LLC, CMT), Ham’s F12 (A549), or RPMI 1640 (H2122) media supplemented with 10% fetal bovine serum, 2mM glutamine, and 100U/ml of penicillin-streptomycin. Platinum-E cells were obtained from Cell BioLabs and used to produce lentivirus encoding Cas9, sgRNA, CD47, and ERK2 by co - transfection of transgene expression vectors with psPAX2 and pMD2.G (Addgene #12260, #12259) using Lipofectamine 3000 (Life Technologies, #L3000015). All cell lines were confirmed to be negative for mycoplasma using a PCR-based assay and were authenticated using short-tandem repeat profiling within the past 2 years.

### 2.2 Generation of CD47 knockout (KO) lines

Clonal CD47 KO lines were generated using CRISPR-Cas9 ribonucleoproteins (RNP) consisting of recombinant Alt-R HiFi S.p. Cas9 nuclease, Alt-R CRISPR-Cas9 tracrRNA, and crRNA targeting CD47 (Integrated DNA Technologies). RNP were delivered to cells using the Neon electroporation system (Thermo Scientific). sgRNA sequences were as follows: hs.sgCD47: AGTGATGCTGTCTCACACAC, and mm.sg.Cd47 TATAGAGCTGAAAAACCGCA. Electroporated cells were clonally expanded and screened for indels within the targeted CD47 coding sequence (CDS) using Sanger sequencing of PCR products spanning the sgRNA cut site. Cells with polyclonal CD47 KO were generated by infecting A549 Cas9+ cells with Lenti-Guide-puro-NLS-GFP (Addgene #185473) encoding an sgRNA targeting human CD47 (sgCD47-6: TAACAGAATTAACCAGAGA). Transduced cells with CD47 perturbation were selected with puromycin and indels were confirmed by PCR and Sanger sequencing. TIDE analysis was used to deconvolute CRISPR-mediated indels generated [25]. Loss of CD47 expression was confirmed as described below. All primers and sgRNA sequences are listed in **Table S1**.

### 2.3 CD47 and ERK2 overexpression

Mouse and human CD47 CDS were PCR amplified from LLC cDNA and a human Mammalian Gene Collection clone purchased from The Centre for Applied Genomics SPARC cDNA Archive, respectively. The CDS for human ERK2 was a generous gift from Dr. William Lockwood and mouse ERK2 was also amplified from LLC cDNA. Amplified CDS were TOPO cloned into pCR8/GW/TOPO (Invitrogen, #K250020). All primer sequences are listed in **Table S1**. A Gateway LR Clonase recombination reaction (Invitrogen, #11791020) was performed to insert the CDS into doxycycline(Dox)-inducible lentivectors: pInducer20-Blasticidin-Neomycin (Addgene #109334) for mouse or pInducer20-Neomycin for human (Addgene #44012). Cloned CDS were confirmed with Sanger sequencing. Lentivirus was produced as described above and stable lines were selected and maintained in media prepared with tetracycline-free FBS containing 4 ug/mL blasticidin or 0.2 mg/ml neomycin. Dox-induced expression of CD47 or ERK transgenes in pInducer20 cells was achieved by 48-72hr of treatment with 100-1000ng/ml Dox and confirmed by flow cytometry as described below.

### 2.4 Immunofluorescence, immunoblotting, flow cytometry and real-time quantitative PCR

Immunofluorescence (IF) was used to detect CD47 expression in cells fixed with 2% paraformaldehyde using the following antibodies: anti-mouse Cd47 (1:1000, Biolegend #127514), anti-human CD47 (1:1000, Biolegend #323124), anti-goat AF555 (1:1500, Thermo Fisher Scientific) and anti-mouse AF488 (1:1500, Thermo Fisher Scientific). DNA was stained with DAPI. Western blots were done using the following antibodies on lysates prepared with RIPA buffer from serum-starved cells: mouse CD47 (1:1000, R&D Systems AF1866), human CD47 (1:1000, Cell Signaling Technologies #63000), p44/42 MAPK (Erk1/2) (1:1000, Cell Signaling Technologies #4696S), phospho-p44/42 MAPK (Erk1/2) (1:1000, Cell Signaling Technologies #4370S), Cdc42 (1:1000, Cell Signaling Technologies #2462), pSRC (1:1000, Cell Signaling Technologies #2101), SRC (1:1000, Cell Signaling Technologies #2110), Vimentin (1:1000, Cell Signaling Technologies #5741), N-Cadherin (1:1000, Cell Signaling Technologies #13116), E-cadherin (1:1000, Cell Signaling Technologies #3195), GAPDH (1:5000, Cell Signaling Technologies #97166S), Vinculin (Cell Signaling Technologies #13901S). Anti-mouse IRDye 800CW and anti-rabbit IRDye 680RD secondary antibodies were purchased from LiCORbio and used at 1:10,000 dilution. Fluorescence imaging of western blots was performed using the LiCOR Odyssey imaging system. Band intensities were background subtracted and normalized to loading controls using ImageStudio software. For flow cytometry, cultured cells were lifted with TrypLE and washed in FACs buffer (10% BSA in PBS -/- with 2mM EDTA). Cells were stained with DAPI for viability and anti-human CD47 (1:100, Biolegend #323124 or #323106) or anti-mouse Cd47 (1:100, Biolegend #127514 or or #127509) antibodies. Stained cells were analyzed on a Sony SP6800 Spectral Cytometer or a Cytoflex-LX. FlowJo software was used for quantitation. RT-qPCR to measure gene expression levels was performed using PowerTrack SYBR green mastermix (Thermo Fisher Scientific, #A46109) with *HPRT1* as the endogenous control and analyzed using the ΔΔCt method. All qPCR primers are listed in **Table S2.**

### 2.5 RNA-sequencing (RNA-seq) and expression analysis

Total RNA was extracted from cultured LLC WT and CD47 KO cells using the RNeasy Plus Mini Kit (Qiagen, #74104) followed by DNAse treatment. RNA integrity was assessed using the 2100 Bioanalyzer (Agilent). Library preparation (NEB Ultra II Directional polyA mRNA) and paired-end sequencing was done at The Centre for Applied Genomics (Toronto, Canada) generating 30-40 million reads per sample on an Illumina Novaseq. RNA from 3 independent biological replicates for each genotype was sequenced. Quality control for the raw sequence data was done using FastQC. Trimmomatic was used to trim adapter sequences and filter out low-quality reads [26]. Cleaned reads were aligned to the Mus musculus GRCm39 transcriptome using the STAR aligner [27]. Principal component analysis and differential expression analysis were conducted using the DESeq2 package in R [28]. Genes with an absolute log_2_ fold change > 1 and a false-discovery rate (FDR) < 0.05 were considered significantly differentially expressed. Gene set enrichment analysis (GSEA) was performed using ClusterProfiler in R to discover gene expression programs associated with CD47 expression [29] and additional pathway enrichment analyses were done using EnrichR [30].

Normalized RNA-seq data for 135 NSCLC cell lines and 515 lung adenocarcinoma (LUAD) tumors from the The Cancer Genome Atlas cohort were accessed from the Dependency Map (DepMap) Portal (https://depmap.org/portal/) and the National Cancer Institute’s Genomic Data Commons portal, respectively [31,32]. NSCLC cell lines were categorized into tertiles based on CD47 expression and single sample gene set enrichment analysis (ssGSEA) was performed on models with CD47 ranking in the top and bottom tertiles of expression (N=45 each) using the Broad’s GenePattern platform [33]. ssGSEA in GenePattern was also performed on LUAD tumors with CD47 expression ranking in the top and bottom 10% of tumors (N=51 each). ssGSEA scores for signatures of EMT and MAPK/ERK signaling (**Table S3**) were extracted and statistically compared between cell lines and tumors with low versus high CD47 expression to assess whether molecular signatures associated with CD47 expression in isogenic LLC cells were also associated with CD47 in independent cell lines and tumors from patients.

### 2.6 Proliferation, and multi-colour competition assays

For proliferation assays, cells were plated at ∼10% confluence in a 24-well dish and images were captured every 2 hours over 3-4 days using the BioTek BioSpa live cell imaging system coupled to a Cytation 5 multimode imaging reader (Agilent). BioTek Gen 5 software was used to calculate percent confluence (i.e., surface area covered by cells) for each well over the course of the experiment. As an orthogonal approach to assess the effect of CD47 KO on cell proliferation, we performed multi-colour competition assays. LLC, CMT, and H1299 cells were engineered with stable Cas9 expression using Lenti-Cas9-blast (Addgene #52962). sgRNA targeting AAVS1 (human negative control), LacZ (mouse negative control), or CD47 were cloned into Lenti-Guide-puro-NLS-GFP or LentiGuide-puro-NLS-mCherry (Addgene #185473 and #185474) and transduced into Cas9+ cells. sgRNA sequences are provided in **Table S4**. mCherry+ sgAAVS1 or sgLacZ were mixed 1:1 with GFP + sgCD47/sgAAVS1/sgLacz cells to compare the relative fitness of cells with and without CD47 perturbation. The ratio of GFP+ to mCherry+ cells was measured over multiple passages by flow cytometry.

### 2.7 Drug dose response assays

Response of cell lines to paclitaxel and cisplatin was evaluated using dose–response assays. Briefly, cells were seeded into 96-well plates and treated with 9 serial dilutions of each drug or DMSO as a control. After 5 days of treatment, cells were fixed with 10% trichloroacetic acid (TCA), stained with sulforhodamine B (SRB), solubilized with 10 mM Tris⋅HCl, and absorbance at 570nm was quantified on a spectrophotometer as a readout of cell viability. Relative viability was calculated as the proportion of SRB signal in drug-treated wells relative to DMSO-treated wells. Dose-response curves were plotted and IC_50_ values were calculated in GraphPad.

### 2.8 Cell adhesion and scratch migration assays

For adhesion assays, 96-well plates were coated with 1 mg/mL fibronectin (Sigma-Aldrich, #F0895) for 30 minutes at room temperature prior to plating cells. Fibronectin was selected as a substrate because it is a component of the extracellular matrix that facilitates EMT processes and CD47 has been shown to mediate binding of reticulocytes and endothelial cells to fibronectin coated surfaces [34–36]. LLC and H1299 cells were seeded onto fibronectin-coated plates and left to adhere at 37°C for 60 minutes. Plates were washed with PBS three times to remove unadhered cells. Adherent cells were fixed in 10% TCA and stained with SRB, which was then solubilized and quantified as absorbance at 570nm on a spectrophotometer (Molecular Devices). Scratch migration assays were done in 24-well plates coated with 1 mg/mL fibronectin. Cells were plated to reach 80-90% confluence 24 hours later. Confluent monolayers were scratched using the BioTek Autoscratch wound making tool. Plates were imaged using the BioTek BioSpa and Cytation 5 live cell imaging system (Agilent) every 2-4 hours until wound closure. Scratch areas were analyzed using the BioTek Gen 5 software.

### 2.9 Tail vein metastasis studies

All animal procedures were performed in accordance with our institutionally approved animal use protocol and the guidelines of the Canadian Council on Animal Care. NSCLC cells were injected into the lateral caudal vein (tail vein) of mice. LLC cells (1 x 10^5^) were injected into syngeneic C57BL/6 mice while H1299 cells (4 x 10^5^) were injected into Nude/Nude mice (N=8 per genotype). We leveraged the luciferase reporter expressed in LLC cells to monitor mice for the development of lung metastases in real-time with bioluminescence imaging done every 2-3 days. Briefly, mice were intraperitoneally injected with 2.25mg luciferin (Revvity, #122799) and imaged on a Newton 7.0 FT-500 Bioluminescent Animal Imager (Scintica) over a 30 minute window to detect peak luminescence signal. A background subtracted radiance threshold of 1x10^5^ p/s/cm^2^/sr was used to define the presence of LLC lung metastases. Mice were euthanized at humane endpoints associated with metastatic tumor burden in the lungs, including mouth breathing, ruffled coat, and hunched posture. Metastasis-free (LLC) and overall (H1299) survival curves were compared between mice injected with WT and CD47 KO cells. Lung tissues were harvested from mice at humane endpoints, formalin-fixed, paraffin-embedded, and tissue sections were used for hematoxylin and eosin staining.

### 3.0 Statistical analysis

The statistical tests used for data analyses are described in the figure legends. All statistics and plots were generated using R and GraphPad Prism software. For cell culture assays, 3 -9 independent biological replicates were analyzed, each with 3-5 technical replicates done to ensure reproducibility of results. Unpaired, two-tailed student’s t-tests were done on continuous data from independent experimental replicates assumed to be random representative samples from a normally distributed dataset with equal variance. One-sample t-tests were used to compare fold-change data setting the hypothetical value to 1 for control samples. Kaplan Meier Survival curves were compared using a log-rank test. ssGSEA scores were compared between cell lines and tumors with low versus high *CD47* mRNA expression using Mann Whitney tests. For all statistical tests, p < 0.05 was considered significant with *p < 0.05, **p < 0.01, ***p < 0.001, and ****p < 0.0001. All error bars in the figures indicate mean ± standard deviation.

## 3.0 Results

### 3.1 Establishment of NSCLC models with CD47 loss of function

To investigate cancer cell-intrinsic signaling and phenotypes influenced by CD47, we selected a panel of 4 widely-used murine and human cell lines with phenotypic features representative of NSCLC seen in patients: murine Lewis lung carcinoma (LLC; Nras- and Kras-mutant; mesenchymal-like) and CMT167 (CMT; Kras-mutant; epithelial-like), as well as human NCI-H1299 (H1299; NRAS-mutant; mesenchymal-like) and A549 (KRAS-mutant; epithelial-like). We used CRISPR to engineer derivative lines lacking the transmembrane and extracellular domains of CD47, which localize CD47 to the plasma membrane and allow it to bind to extracellular ligands (**Fig.1A)**. Sanger sequencing of LLC, CMT, and H1299 single cell-derived clonal KO lines and A549 cells with polyclonal editing validated the presence of CRISPR-induced indels in the *CD47* gene **(Fig.S1A)**. These indels caused frameshifts resulting in truncations of CD47 protein **(Fig.1A)**. All KO lines were viable. Loss of CD47 expression in these lines was validated by quantitative real-time PCR and flow cytometry **(Fig.1B-C)**, as well as immunoblotting and immunofluorescence (**Fig.S1B-C**), confirming their utility for studying CD47-mediated signaling and its associated phenotypic effects in NSCLC cells.

**Figure 1.**
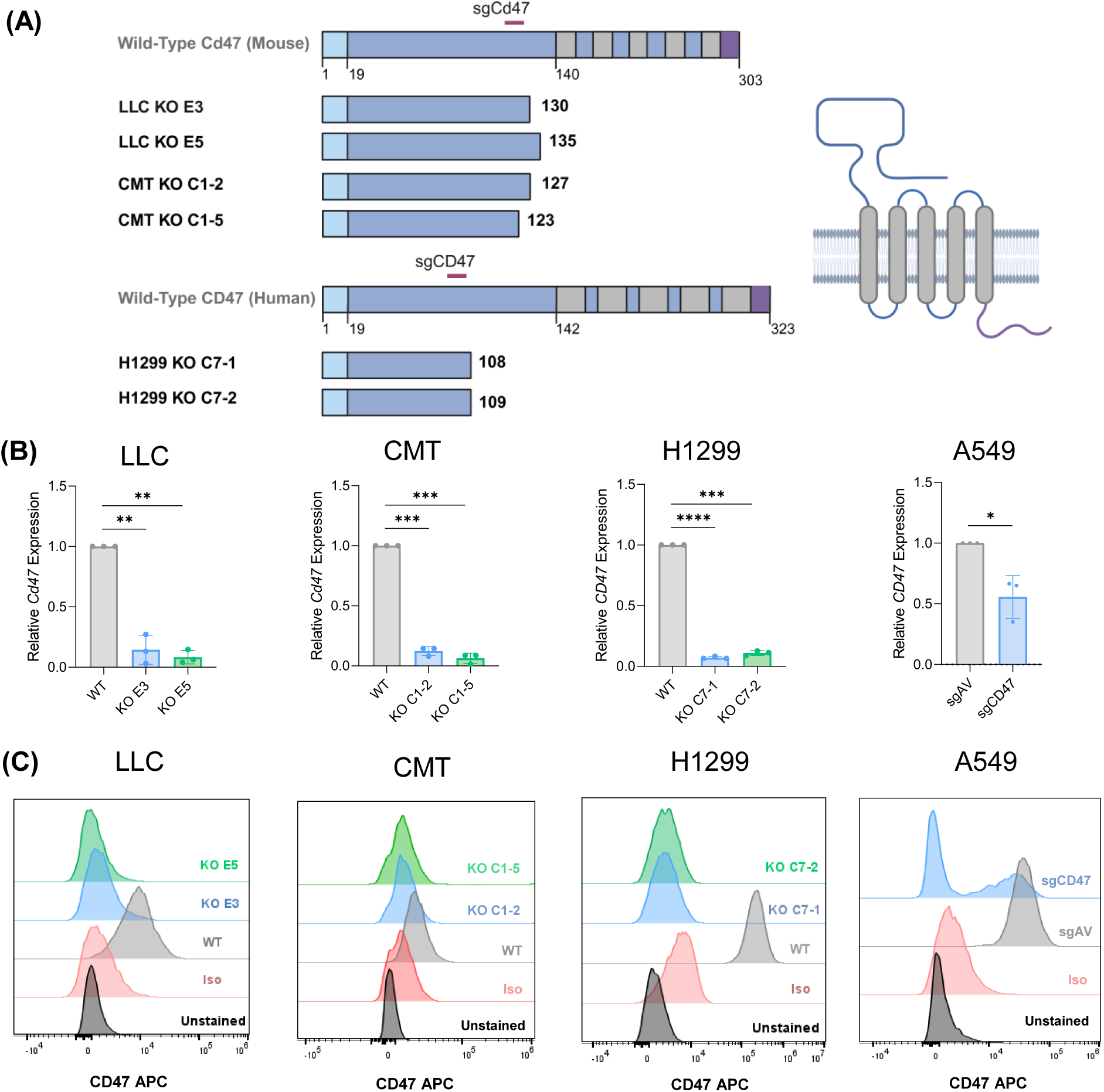
Generation of isogenic NSCLC models with loss of CD47 expression. **A)** Schematic representation of LLC, CMT, and H1299 cell lines engineered with deletion of the C-terminal half of CD47, which encodes its transmembrane and cytoplasmic tail domains. The signal peptide (light blue), transmembrane domains (grey), extra/intracellular domains (dark blue), and cytoplasmic tail (purple) are indicated. The protein structure for CD47 was obtained from UnitProt. CD47 knockout (KO) lines were established following electroporation of cells with Cas9/sgRNA ribonucleoproteins followed by clonal expansion and characterization. A549 cells with polyclonal knockdown of CD47 were also generated by infecting A549-Cas9+ cells with sgRNA targeting CD47 (sgCD47) or AAVS1 as a negative control (sgAV). Transduced sgAV and sgCD47 cells were selected with puromycin. In all models generated, loss of CD47 was verified by qPCR for mRNA expression **(B)** and flow cytometry for protein expression on the cell surface **(C)**. Fold-changes in mRNA expression of CD47 in WT and KO cells were statistically assessed using one-sample t-tests. For cytometry, unstained cells and cells stained with an isotype control antibody (Iso) are also shown. CRISPR-induced indels in the *CD47* gene were also verified by Sanger sequencing, and loss or protein expression was further confirmed with immunoblotting and immunofluorescence (Fig. S1).

### 3.2 WT and CD47 KO NSCLC cells exhibit distinct gene expression profiles

We employed a transcriptomic approach to discover how CD47 influences gene expression patterns in NSCLC cells. RNA-seq data was generated for our LLC lines as part of a different study by our group (unpublished). Gene expression analyses of these isogenic models revealed that WT and CD47 KO cells were transcriptionally distinct as they clustered separately in principal component analysis, with CD47 among the genes segregating the principal components (**Fig.2A;Fig.S2A,C**). Differences in the gene expression profiles of WT and KO cells were apparent upon visualizing the 500 most variably expressed genes (**Fig.2B)**. We next performed comparative analyses to discover genes differentially expressed between WT and KO cells for each clone. This identified 4,875 differentially expressed genes (DEGs) in KO clone E3 and 3,567 in KO clone E5 (FDR < 0.05). Comparing these lists revealed 1,842 DEGs common to both clones (**Fig.2C)**. We then separated the DEG lists into up- and downregulated genes, defined as having a log_2_ fold-change in expression >1 or <-1 in KO versus WT cells, respectively, and compared them across the clones. This revealed 1,450 DEGs with matching directions in both clones. As expected, CD47 was identified as a top downregulated gene in KO relative to WT cells. These results suggest that CD47 regulates specific gene expression networks in NSCLC.

**Figure 2.**
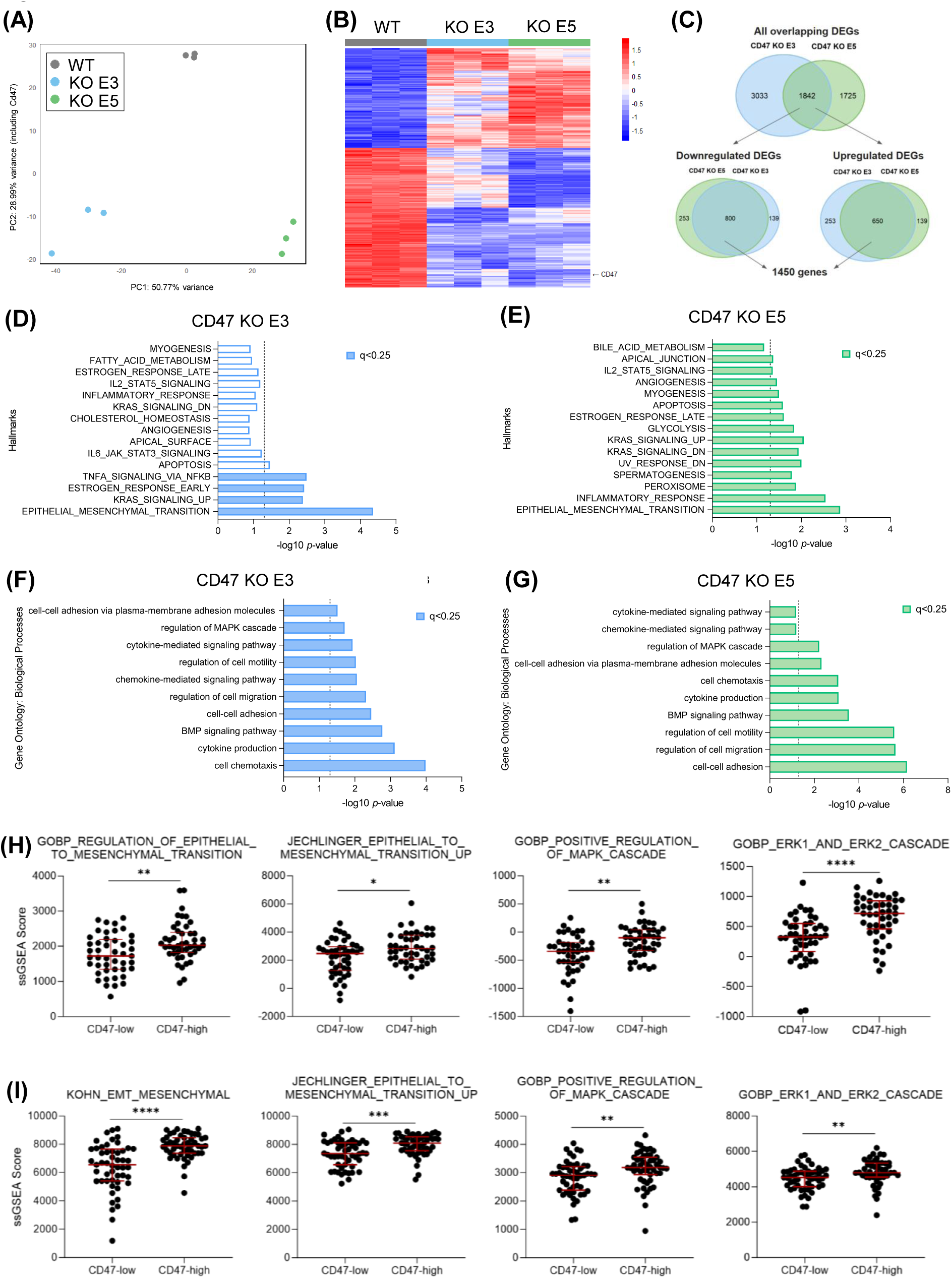
CD47 KO cells exhibit transcriptomic changes associated with downregulation of EMT pathways. **A)** Principal component analysis (PCA) indicating dissimilarity in transcriptomes of LLC WT and CD47 KO cell lines. PCA was performed on gene expression profiles generated by RNA-seq with 3 replicates per line. **B)** Heatmap of the 500 most variably expressed genes in LLC WT and CD47 KO cells, which includes CD47 as one of the most significantly downregulated genes in CD47 KO cells. **C)** Statistically significant differentially expressed genes (DEGs) identified by DEseq2 common to both CD47 KO clones. **D-G)** Gene set enrichment analysis (GSEA) results indicating MSigDB Hallmark and Gene Ontology: Biological Processes gene sets enriched for downregulated DEGs in CD47 KO cells. The dotted vertical line indicates an adjusted p-value < 0.05. **H)** ssGSEA scores for EMT and MAPK gene expression signatures in NSCLC cell line RNA-seq data from DepMap. Models were stratified into tertiles based on CD47 expression (N=45 per group) and ssGSEA scores were compared between cell lines with the lowest and highest CD47 expression (i.e., top and bottom tertiles). **I)** ssGSEA analysis of RNA-seq data for lung adenocarcinoma tumors accessed from the TCGA database. Similar to (H), ssGSEA scores were compared between tumors with the lowest and highest CD47 expression levels (i.e., top and bottom 10%; N=51 per group). Mann Whitney tests were used to compare ssGSEA scores in CD47 high and low samples in (H,I).

### 3.3 CD47 governs MAPK and EMT transcription programs in lung cancer cells

We proceeded to determine what biological pathways are affected by gene expression changes driven by CD47 KO in the LLC model. We reasoned that genes downregulated in KO cells would be enriched in pathways that promote cell-intrinsic phenotypes governed by CD47. Accordingly, GSEA on DEGs downregulated in CD47 KO cells identified “Epithelial Mesenchymal Transition” (EMT) as the most significant Hallmark pathway enriched in both KO clones (**Fig.2D-E**). GSEA of downregulated DEGs considering Biological Processes annotated in the Gene Ontology knowledgebase revealed several EMT-related processes including “chemotaxis”, “cell-cell adhesion”, “cytokine production”, “cell migration” and “regulation of MAPK cascade” as significantly enriched (**Fig.2F-G)**. Further confirming the positive association between CD47 and EMT, analyses using the EnrichR tool indicated that genes downregulated in LLC CD47 KO cells were enriched for targets of the EMT-promoting transcription factors, Twist and Prrx2 **(Fig.S2D)**. Assessment of the individual DEGs driving enrichment of the cell adhesion, cell migration, and MAPK signaling pathways revealed numerous genes encoding glycoproteins, matricellular proteins, ligands, and growth factors that contribute to the extracellular matrix and promote EMT **(Fig.S2E-G)**. These findings indicate for the first time that CD47 influences the transcriptome and positively regulates MAPK and EMT-associated gene expression signatures in NSCLC cells.

To corroborate these findings, we investigated associations between CD47 expression, MAPK and EMT molecular signatures in a panel of 135 NSCLC cell lines using publicly available transcriptomic data from the Broad Institute’s DepMap portal (https://depmap.org/portal/) [31]. We segregated these models into groups based on CD47 expression and performed single sample GSEA (ssGSEA) on cell lines ranking in the top and bottom tertiles of CD47 expression. Consistent with our LLC RNA-seq-derived results, we found that NSCLC cell lines with high CD47 expression had significantly greater ssGSEA scores for gene expression signatures of EMT and MAPK signaling compared to lines with low CD47 expression (**Fig.2H**). To assess the clinical relevance of this finding, we performed the same transcriptomic analysis on lung tumors from the TCGA cohort. We obtained RNA-seq data for 515 lung adenocarcinomas, the most common type of NSCLC, and compared ssGSEA scores between tumors with the highest and lowest expression of CD47 (top and bottom 10 percent of tumors). Consistent with the cell line results, tumors with high CD47 expression exhibited significantly greater enrichment scores for EMT and MAPK pathway signatures (**Fig.2I**). These additional transcriptomic analyses in large cohorts of cell lines and clinically relevant tumor specimens provide further evidence supporting a generalizable role for CD47 in positively regulating MAPK and EMT expression programs in NSCLC.

### 3.4 Loss of CD47 is associated with reduced expression of mesenchymal markers

As an initial step to validate the role of CD47 in promoting EMT in lung cancer, we assessed established EMT markers in our isogenic NSCLC cell lines. LLC and H1299 are inherently mesenchymal-like and are commonly used to model metastasis *in vivo* [37,38], enabling us to assess whether CD47 contributes to these phenotypes in these models. Accordingly, we did not detect E-cadherin in either line (**Fig.3A,C**). In LLC cells, we observed a significant decrease in N-cadherin expression in both CD47 KO clones and a significant reduction in vimentin expression in one of two clones (**Fig.3A**). In H1299 cells, N-cadherin was also significantly downregulated in one KO clone with a non-significant trend evident in the other, and a trend towards reduced vimentin expression was also evident in both clones **(Fig.3C)**. Since the prominent mesenchymal nature of LLC and H1299 cells could limit our ability to detect changes in mesenchymal marker expression, we also tested the effect of CD47 KO in CMT and A549, which are more epithelial in nature. Both of these lines express E-cadherin, CMT also expresses N-cadherin, and A549 expresses both N-cadherin and vimentin **(Fig.3B,D)**. In CMT cells, we observed trends towards reduced vimentin in both clones and significantly increased E-cadherin expression in one of two clones **(Fig.3B)**. Although some of these CMT results were not significant, the direction of changes observed were reproducible across all experimental replicates. In polyclonal A549 cells, expression of both vimentin and N-cadherin was significantly reduced in cells with CD47 perturbation and a trend towards increased E-cadherin expression was also evident **(Fig.3D).** Although the specific changes in EMT markers that occur upon CD47 KO varied across models, likely due to the heterogeneity characteristic of NSCLC, our observations of reduced mesenchymal marker expression upon CD47 KO were evident in all 4 models and are consistent with CD47 having a pro-EMT function in NSCLC as indicated by our transcriptomic data.

**Figure 3.**
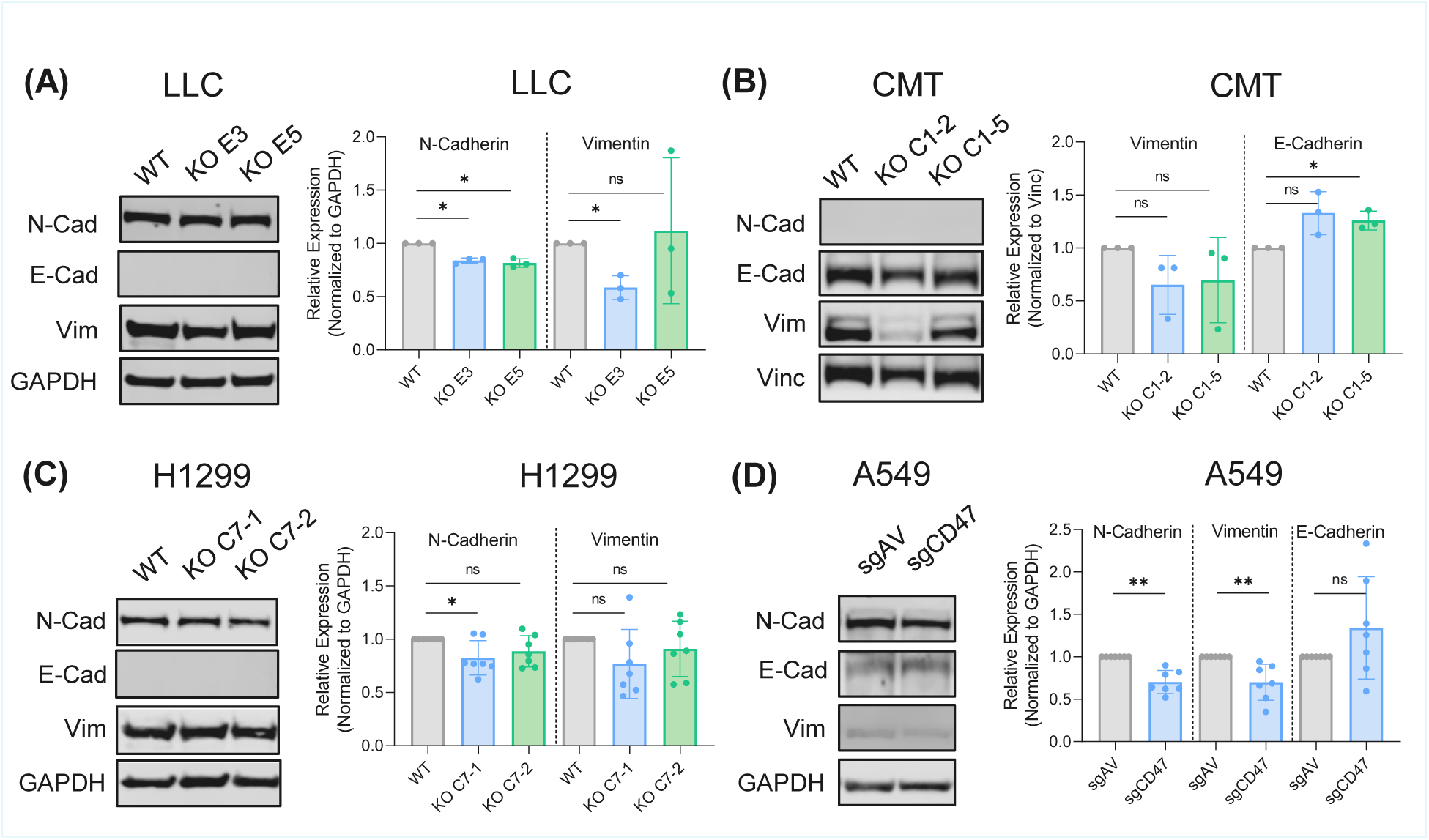
Loss of CD47 expression reduces expression of EMT markers in NSCLC cells. A-D) Expression of the canonical EMT markers E-cadherin, N-cadherin, and Vimentin in NSCLC models isogenic for CD47. LLC and H1299 are mesenchymal-like cells that lack E-cadherin but express vimentin and N-cadherin. CMT and A549 are more epithelial-like, expressing E-cadherin with differential expression of vimentin and N-cadherin. Western blots were conducted on protein lysates from cells that were serum starved for 24 hours. Quantitation of EMT markers was conducted in ImageStudio as described in the methods and the fold change in expression in CD47-deficient relative to control cells was calculated for each marker for each replicate experiment. A one-sample t-test was used to determine whether fold-changes in CD47 KO/knockdown cells differed significantly from the hypothetical value of 1 in WT controls. Error bars indicate mean ± SD.

Since EMT is strongly associated with cancer drug resistance [39], we also tested the sensitivity of WT and KO cells to chemotherapies used to treat NSCLC. We observed no reproducible differences in response to cisplatin across CD47 KO lines (**Fig.S3**). CD47 KO enhanced the sensitivity of LLC and H1299 cells to paclitaxel, evident by significantly lower IC _50_ values in KO compared to WT cells **(Fig.S3A)**. However, loss of CD47 expression did not affect paclitaxel response in CMT and A549, suggesting this phenomenon is not generalizable across NSCLC models **(Fig.S3)**.

### 3.5 CD47 KO impairs EMT-related phenotypes in NSCLC cells

To further assess the intrinsic relevance of CD47 in NSCLC cells and functionally validate our RNA-seq findings, we investigated the impact of CD47 KO on various malignant phenotypes including those characteristic of EMT. We first tested cell proliferation as several studies have reported that CD47 regulates this essential process in other cancer types [16]. However, live cell imaging of LLC, CMT, H1299, and A549 cells to monitor confluence over time revealed that CD47 KO had no effect on proliferation in any model **(Fig.S4A)**. As an orthogonal approach to confirm the lack of proliferation phenotype in CD47-deficient cells seen in other cancer types, we also compared the relative proliferation of LLC, CMT, and H1299 cells with WT CD47 versus CRISPR-mediated CD47 knockdown using multi-colour competition assays **(Fig.S4B)**. We achieved CD47 knockdown using this approach but again observed no difference in the fitness of CD47-deficient and unperturbed control cells over multiple passages **(Fig.S4C-D)**. These results indicate CD47 is dispensable for proliferation in NSCLC cells.

Next, we assessed the effects of CD47 KO on cell adhesion and migration, two processes nominated by our transcriptome analyses that are critical for EMT, cancer metastasis, and progression. In the LLC, H1299, and A549 models, CD47 KO significantly reduced cell adhesion to plates coated with fibronectin, an extracellular matrix protein that facilitates cancer cell invasion and metastasis [36] (**Fig.4A; Fig.S5A**). We observed no effect of CD47 KO on adhesion in CMT cells (**Fig.4A)**. We used scratch wound assays to determine the effects of CD47 KO on cell migration. Time lapse imaging of cell monolayers after the scratch was applied revealed that loss of CD47 expression significantly impaired the ability of cells to migrate in the A549 model, both LLC and CMT KO clones, and one of two KO clones in H1299 (**Fig.4B;Fig.S5B**), consistent with findings in a previous NSCLC study [13]. These experiments indicate that the EMT expression signatures we identified as downregulated in CD47 KO cells and in NSCLC cell lines and tumors with low CD47 expression are associated with corresponding defects in cell adhesion and migration, functionally validating our transcriptomic findings. To confirm the role of CD47 in mediating these phenotypes, we engineered CD47-deficient LLC and H1299 cells with dox-inducible CD47 expression **(Fig.4C)**, as CD47 KO in these lines most strongly suppressed cell adhesion and migration **(Fig.4A-B)**. Re-constitution of CD47 rescued the migration defect seen in LLC KO cells and enhanced cell adhesion in one KO clone with the same trend in the other **(Fig.4D-E)**; re-expression also rescued KO-associated defects in adhesion and migration in the H1299 model **(Fig.4D-E)**. Furthermore, we engineered a human NSCLC cell line, H2122 (KRAS-mutant), with low endogenous CD47 expression, to express the dox-inducible CD47 system. Upregulating CD47 expression in these cells led to a non-significant trend towards enhanced cell adhesion and a significant increase in cell migration **(Fig.4C-E)**. Collectively, these functional experiments indicate that CD47 promotes the adhesive and migratory abilities of NSCLC cells, consistent with a pro-EMT function.

**Figure 4.**
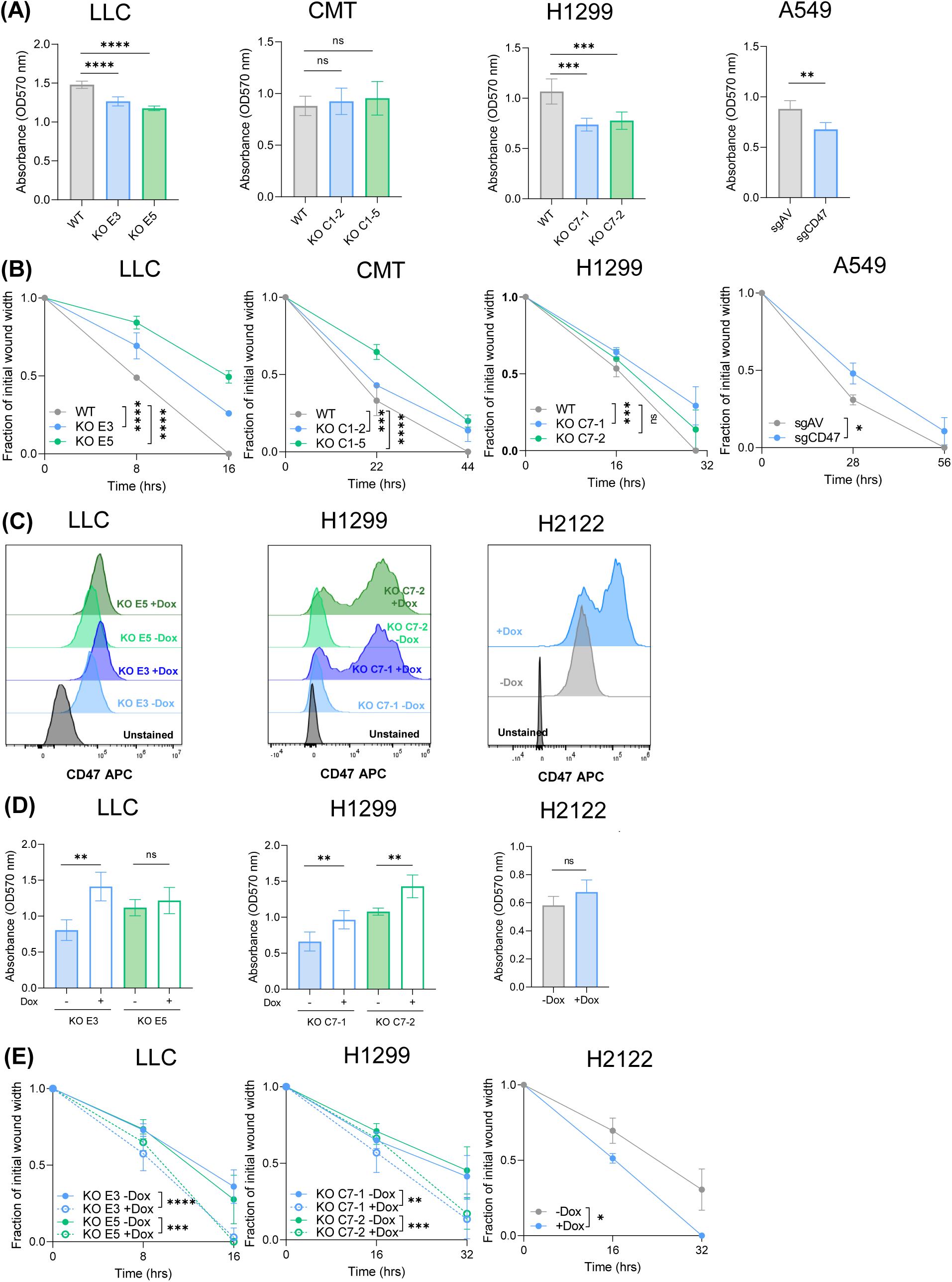
Loss of CD47 impairs lung cancer adhesion and migration. **A)** Cell adhesion assay results in LLC, CMT, H1299 and A549 models. Cells were plated in 96-well plates coated with fibronectin. One hour after seeding, cells were washed with PBS and fixed with TCA, followed by treatment with sulforhodamine B (SRB) to stain adherent cells. SRB absorbance values were compared across genotypes to identify differences in cell adhesion. **B)** Wound healing assays measuring cell migration after scratching a confluent cell monolayer using the BioTek Autoscratch robot. Wound closure was monitored by capturing images of the scratched area every 2-4 hours using the BioTek Cytation 5 live cell imaging instrument. Wound width was quantified and compared across genotypes immediately after the scratch and at two time points after the scratch and normalized to the initial scratch width. Asterisks shown indicate statistical significance at assay endpoints. Data presented for cell adhesion and migration assays are representative of at least 3 independent experiments. **C)** Flow cytometry confirmation of surface level CD47 expression after doxycycline (Dox) induction in LLC and H1299 CD47 KO lines and H2122 cells with low endogenous CD47 expression. Cytometry was performed 48-72hr after treatment with 100-1000ng/ml Dox. **D)** Results for cell adhesion assays in CD47-deficient LLC and H1299 cells and CD47-low H2122 cells engineered with the pInducer20-CD47 system. Cells were treated with and without Dox (100-1000ng/ml) to induce CD47 expression as shown in (C). Adhesion assays results are representative of 3 independent experiments. **E)** Migration assay results for the same cells with CD47 expression induced by dox. Migration assays presented are representative of 3 independent experiments. Dotted lines indicate the +Dox condition in migration assays. Comparisons of cell adhesion and migration between groups were done using two-tailed t-tests.

### 3.6 CD47 positively regulates MAPK signaling to promote EMT

We next sought to determine how CD47 signaling regulates EMT in NSCLC cells. Studies in epithelial cells, neurons, platelets and/or cancer cells identified SRC, FAK, and Cdc42 as downstream mediators of CD47 signaling that stimulate pro-migratory phenotypes [16], but we found no evidence that CD47 regulates these effector proteins in our NSCLC cell lines ( **Fig.S6**). Since our RNA-seq analyses implicated the MAPK cascade as a putative downstream effector of CD47 (**Fig.2**), and MAPK signaling is known to drive EMT [40], we hypothesized that CD47 signals through MAPK to induce pro-EMT transcription programs and associated cellular phenotypes. To test this and confirm our transcriptomic findings, we evaluated the effects of CD47 KO on MAPK signaling using phospho-ERK as a readout of pathway activity. Consistent with our hypothesis, we found that phospho-ERK was reduced in LLC, H1299, and CMT cells with CD47 KO compared to WT controls (**Fig.5A-C)**, but no significant difference in phospho-ERK expression was seen in A549 cells with polyclonal CD47 knockdown (**Fig.5D)**. This could reflect incomplete KO of CD47 in this model or suggest that ERK is not an effector of CD47 in A549 cells. Nevertheless, consistent with ERK being a downstream mediator of CD47 signaling in NSCLC cells, we observed that re-expression of CD47 tended to increase phospho-ERK levels in CD47-deficient LLC cells and significantly increased phospho-ERK in H1299 KO lines, both of which exhibited strong downregulation of phospho-ERK **(Fig.5E-F)**. Furthermore, we observed a non-significant trend towards increased phospho-ERK expression in CD47-low H2122 cells upon dox-induced expression of CD47 **(Fig.5G)**.

**Figure 5.**
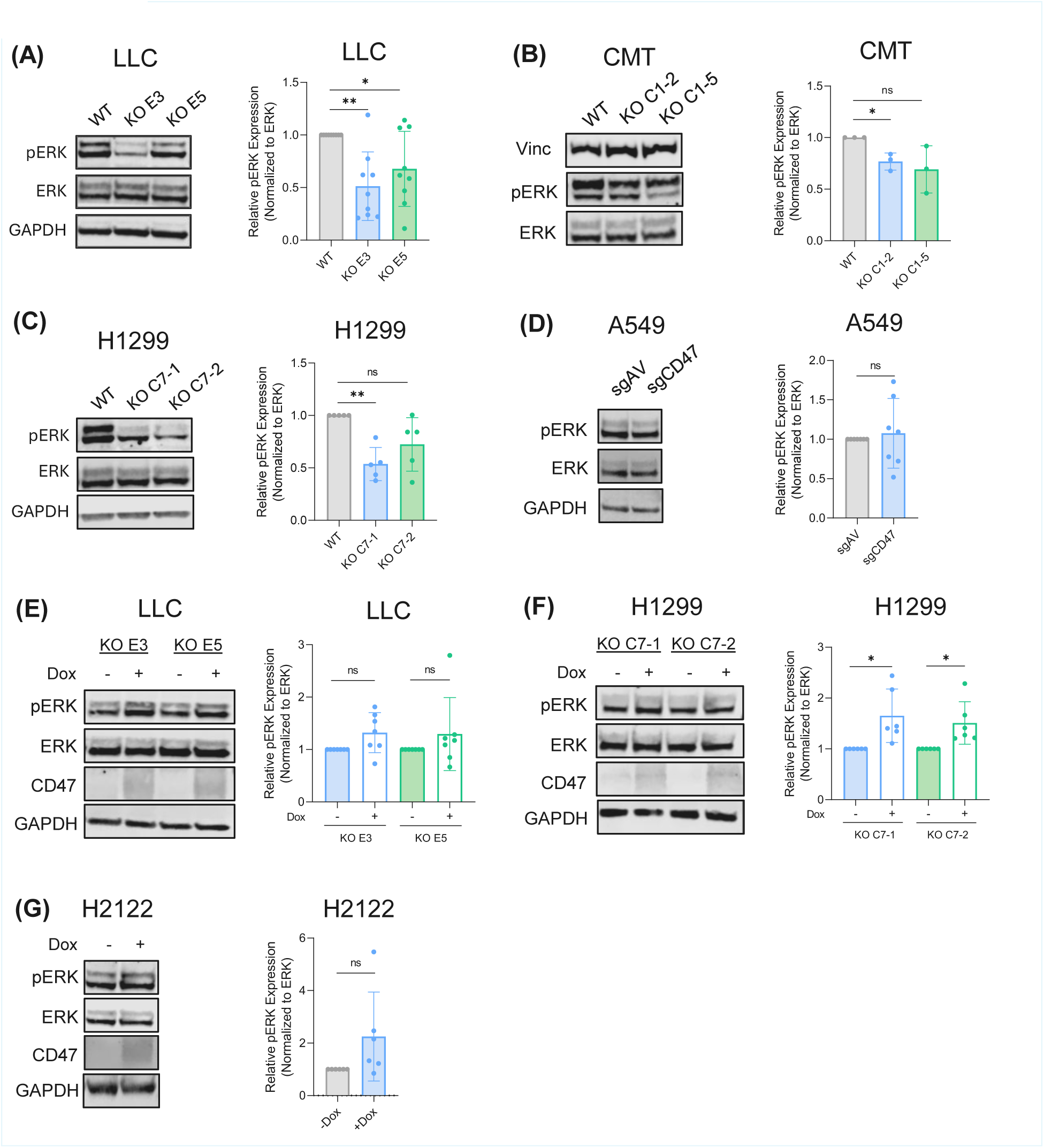
CD47 regulates MAPK signaling in lung cancer cells. A-D) Western blots and quantitation of phospho-ERK (pERK) expression in WT versus CD47 KO or knockdown cells serum starved for 24 hours. pERK expression was normalized to total ERK. pERK expression is presented as the fold-change in CD47 KO/knockdown cells relative to unperturbed wildtype controls, and compared across genotypes using a one-sample t-test. **E-G)** Immunoblotting for CD47 and pERK in CD47-deficient LLC and H1299 cells and H2122 cells with low endogenous CD47 expression treated with and without Dox (100-1000ng/ml) to induce CD47 expression. The fold-change in pERK expression between cells deficient for CD47 and reconstituted with CD47 is illustrated and differences in pERK fold-change were statistically compared using a one-sample t-test. Each data point indicates an individual biological replicate.

We then investigated whether elevating ERK expression in CD47-deficient cells also reversed defects in cell adhesion and migration using the same dox-inducible system. We selected LLC and H1299 for these studies since CD47 KO had the largest magnitude effect on reducing phospho-ERK in these models. We were unable to robustly induce ERK expression in the LLC KO E3 clone but we achieved reproducible increases in ERK expression in clone E5 and both H1299 KO clones **(Fig.6A-B)**. Despite all experimental replicates exhibiting ERK overexpression in H1299 cells, the fold-change in ERK expression was not significantly different in cells treated with and without dox. However, increasing ERK expression rescued cell adhesion and migration deficiencies in CD47 KO LLC cells **(Fig.6C-D),** rescued cell adhesion in both H1299 KO clones **(Fig.6E),** and enhanced cell migration in one H1299 KO clone with a reproducible trend observed in the other **(Fig.6F)**. Taken together, these data suggest that CD47-mediated regulation of EMT phenotypes is dependent on MAPK signaling, at least in these NSCLC models.

**Figure 6.**
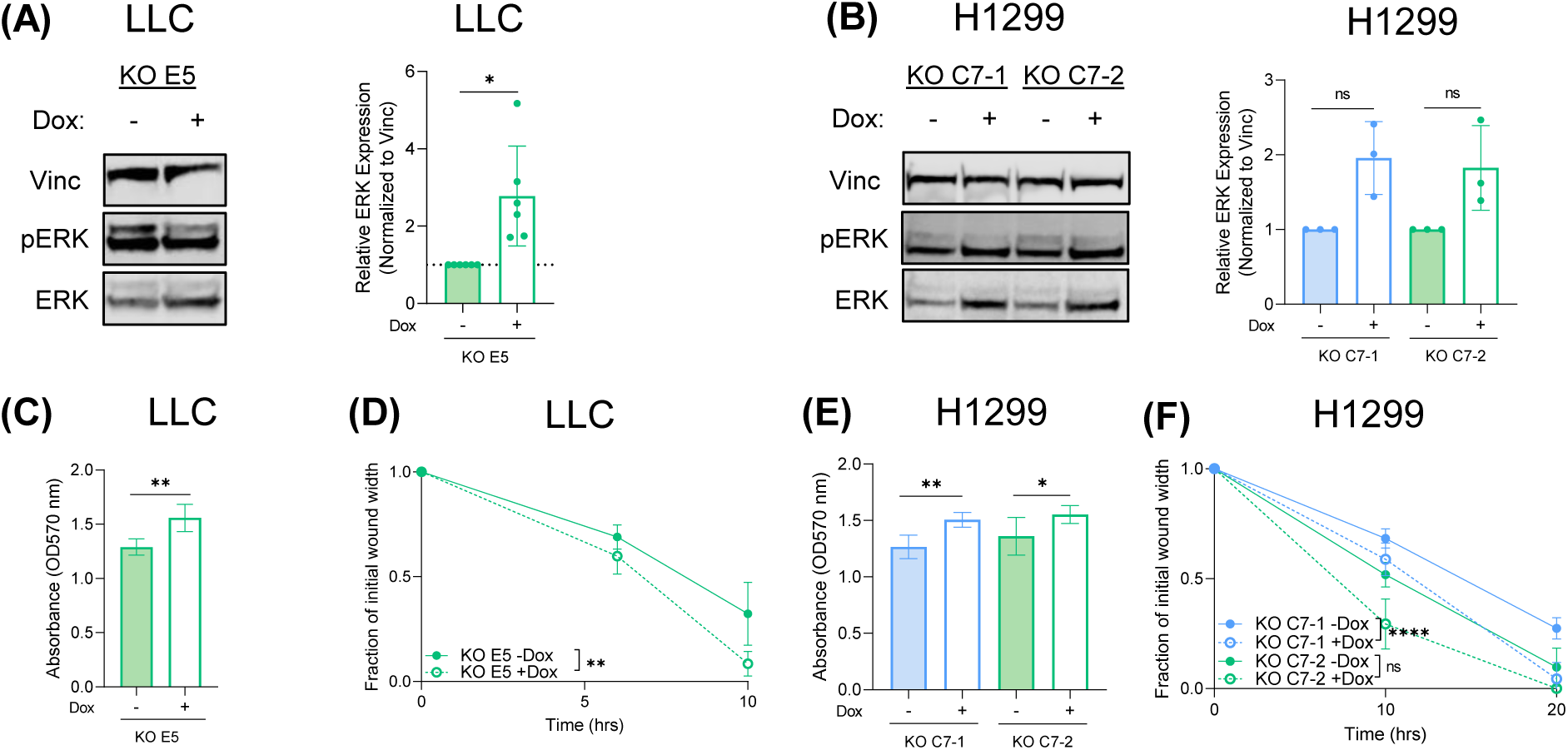
CD47-mediated regulation of cell adhesion and migration is ERK dependent. A,B) Immunoblots and quantitation confirming phospho-ERK (pERK) upregulation in LLC and H1299 WT and CD47 KO cells expressing a dox-inducible ERK2 construct treated with and without 500 ng/ml (LLC) or 100ng/ml (H1299) Dox for 24-48h. pERK expression is normalized to Vinculin (Vinc) since dox treatment induces the expression of total ERK. Fold-changes in ERK expression were compared between -/+ Dox conditions using a one-sample t-test. **C-F)** Results for cell adhesion and migration assays in the same ERK-inducible LLC and H1299 cells shown in (A,B) pre-treated with or without 100-500ng/ml Dox for 24-48h. Asterisks shown indicate statistical significance for two-tailed t-tests. Data presented are representative of at least 3 replicate experiments. Dotted lines indicate the +Dox condition in migration assays.

### 3.7 CD47 KO diminishes NSCLC metastasis in vivo

Finally, we assessed the physiological relevance of the CD47-ERK-EMT signaling axis we discovered using an intravenous model of metastasis in mice that models cancer cell survival in circulation, cell seeding at a distant site, and metastatic tumor outgrowth. We assessed metastasis of LLC and H1299 cells because they have previously been reported to metastasize to the lungs after intravenous injection into mice. We injected LLC and H1299 WT and CD47 KO cells into the tail vein of syngeneic C57BL/6 mice and immunocompromised Nude/Nude mice, respectively **(Fig.7A)**. Mice were euthanized at humane endpoints associated with tumor burden. In both models, we found that CD47-deficient cells had a dramatically impaired ability to form metastatic tumors in the lungs, which was evident from bioluminescence imaging for LLC **(Fig.7B, Fig.S7)** and inspecting the lungs and thoracic cavity of mice at necropsy for H1299 **(Fig.7F)**. The formation of tumor nodules in C57BL/6 mice injected with CD47 KO LLC cells indicates their ability to evade phagocytes and engraft in the lungs **(Fig.7D)**. Neither tumor nodules nor micro metastases were evident in the lungs of mice injected with CD47-deficient H1299 cells **(Fig.7F)**, although some Nude/Nude mice in the CD47 KO group were sacrificed due to hind limb paralysis and significant body weight loss. This could suggest that CD47 KO cells engrafted to form extra-pulmonary metastases elsewhere, and/or that phagocyte-mediated clearance of CD47 KO cells contributed to the lack of lung metastases in the H1299 model. For both models, mice injected with CD47 KO cells exhibited significantly longer metastasis-free and overall survival times than control mice injected with WT cells **(Fig.7C,E)**. These results suggest that CD47 and the cell-intrinsic MAPK-EMT program it regulates supports NSCLC metastasis *in vivo*, providing evidence of its physiological relevance to tumor progression.

**Figure 7.**
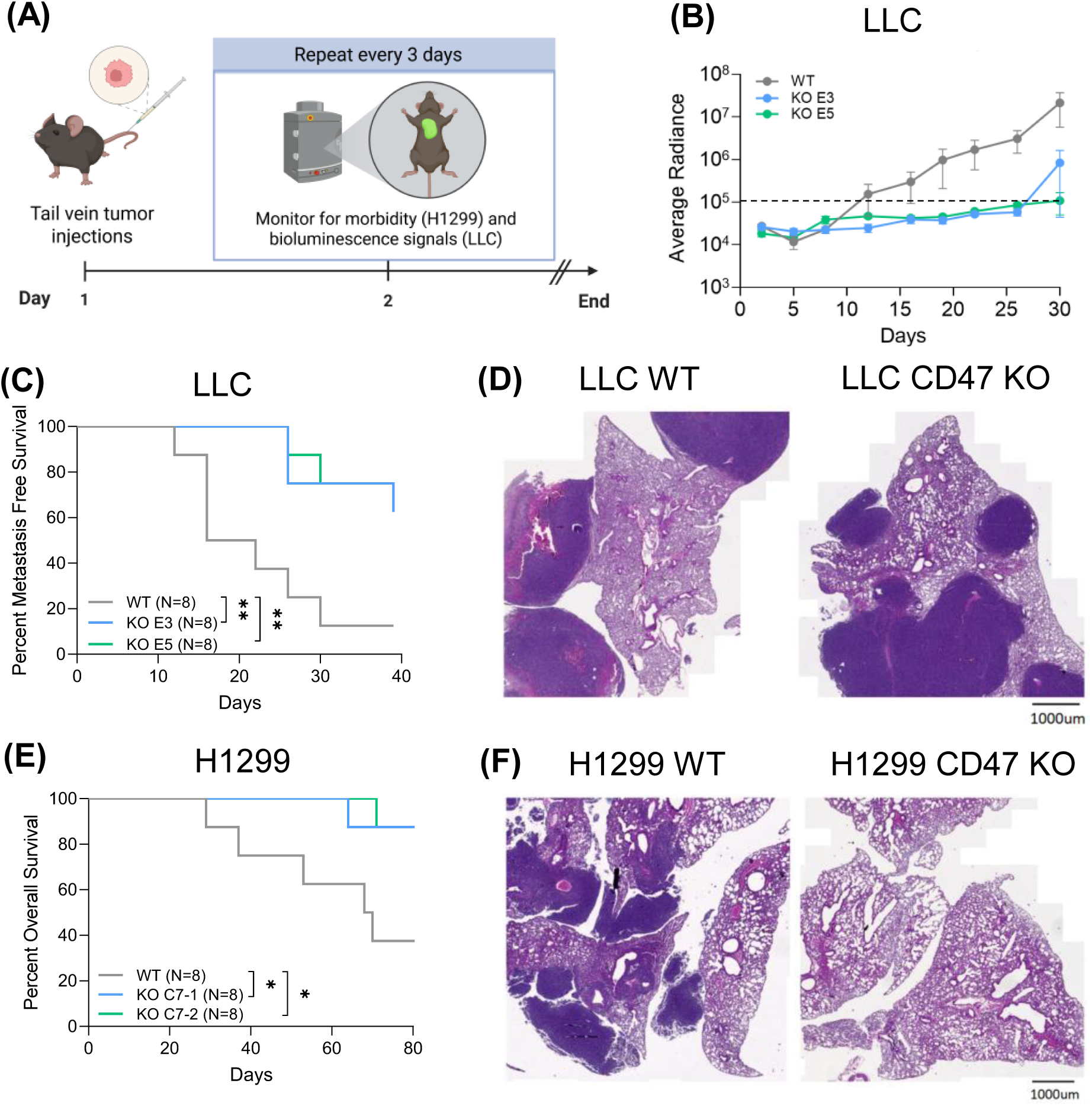
CD47 KO significantly diminishes lung cancer metastasis *in vivo*. **A)** Schematic indicating the experimental design for assays evaluating the metastatic abilities of wildtype and CD47 KO LLC and H1299 cells in mice. Cells were delivered via injection into the tail vein of syngeneic C57BL/6 (LLC) or immunodeficient Nude/Nude (H1299) mice. Bioluminescence imaging (BLI) after intraperitoneal injection of luciferin was used to detect the emergence of lung tumor nodules for the LLC model which expresses luciferase. Mice injected with H1299 cells were monitored for signs of metastasis-associated morbidity including mouth breathing, ruffled coat, and hunched posture. **B)** Growth curves for LLC-derived lung tumor nodules generated from BLI imaging. Dotted line indicates the radiance signal threshold for defining the presence of lung metastases. **C)** Metastasis-free survival curves for mice injected with LLC wildtype and CD47 KO cells (N=8 per group). **D)** Hematoxylin and eosin (H&E) staining of lung tissues harvested from C57BL/6 mice at endpoint, illustrating lung tumor nodules formed by wildtype and CD47 KO LLC cells. **E)** Overall survival curves for mice injected with H1299 wildtype and CD47 KO cells (N=8 per group). **F)** H&E staining for lungs harvested from Nude/Nude mice injected with H1299 cells at study endpoint. Kaplan-Meier survival curves were compared between mice injected with wildtype and CD47 KO cells using a log-rank test.

## 4.0 Discussion

In this study, we sought to elucidate the cell-autonomous functions of CD47 and define its effects on intracellular signaling and molecular programs specifically in NSCLC. We discovered for the first time that lung cancer cells with CD47 KO have distinct transcriptional profiles characterized by diminished expression of MAPK and EMT pathway signatures. Our functional studies validated these molecular findings, suggesting that CD47 positively regulates EMT phenotypes through an ERK-dependent mechanism in NSCLC. We also found that CD47 LOF significantly diminished metastasis of NSCLC cells *in vivo,* indicating the pathophysiological relevance of the CD47–ERK–EMT axis we discovered and highlighting the potential utility of CD47 blockade for antagonizing tumor metastasis and disease progression in NSCLC patients.

CD47 has been shown to regulate several biological processes that contribute to cellular plasticity and tissue homeostasis, many of which are exploited by cancer cells to promote tumor growth and progression [16]. As recently reviewed by Polara et al., these cell-intrinsic CD47 functions include regulation of proliferation, cell death and survival, metabolism, differentiation, motility, autophagy, senescence and stemness. These functions have been characterized in different cell types including non-malignant glial, epithelial, endothelial, immune, and smooth muscle cells, as well as in various cancer models, and are dictated by the proteins CD47 interacts with [16]. For example, CD47 signaling induced by its ligand TSP-1 stimulates proliferation in astrocytoma and glioblastoma cancer cells but suppresses proliferation in endothelial, NK and T cells, demonstrating the cell-type specificity of CD47’s effects [41–44]. The context-specific functions of CD47 motivated us to investigate cell-intrinsic signaling and phenotypes regulated by CD47 in NSCLC. We observed that CD47 KO had no effect on cell proliferation in NSCLC models, consistent with observations in CD47-deficient small-cell lung cancer cells [15]. However, we found that CD47 KO impaired NSCLC migration and adhesion, two phenotypes associated with EMT and integral for cancer cell metastasis [45]. Unlike its effects on proliferation, positive regulation of cell migration and adhesion by CD47 appears to be common across cell types with this phenotype evident in erythrocytes, neutrophils, smooth muscle, epithelial, melanoma, ovarian, lymphoma, and colorectal cancer cells [16]. The ability of CD47 to promote these phenotypes has been attributed to interactions with TSP-1 and/or integrins in some but not all of these cellular contexts [16].

Our findings advance upon those of a previous study that implicated CD47 in NSCLC invasion and metastasis [13]. Zhao et al. characterized the cellular effects of CD47 using RNA interference (RNAi) and transient CD47 expression in the A549 and NCI-H520 models. They reported that CD47 promotes migration and invasion and has no effect on cell viability, consistent with our findings. This group identified the Rho GTPase, Cdc42, as a downstream effector of CD47-mediated regulation of these phenotypes. However, we observed no change in Cdc42 expression upon CD47 KO in LLC, CMT, H1299 or A549 cells, suggesting Cdc42 may not be a generalizable mediator of CD47 signaling in NSCLC. Given the established role of Cdc42 as an EMT effector [46], Zhao et al. assessed canonical EMT markers in their cell models but found that perturbing CD47 did not alter expression of E-cadherin, N-cadherin, Snail or Slug, leading them to conclude that CD47 does not regulate EMT. While we could not detect expression of the canonical EMT transcription factors, Twist, Snail, or Slug in our models (data not shown), we observed reduced N-cadherin and vimentin expression and trends towards increased E-cadherin expression across multiple NSCLC models with CD47 KO. Although the specific EMT markers whose expression changed upon CD47 KO varied by cell line, every model exhibited an EMT-associated change in at least one marker. Additionally, our transcriptomic data in isogenic NSCLC cells, as well as NSCLC models and patient tumors with low and high CD47 expression, strongly implicated CD47 in regulating EMT at the molecular level. We suspect that our conclusions regarding the role of CD47 in EMT in NSCLC differ from those of Zhao et al. due to model-specific differences and/or differences in the experimental approaches used. With respect to the former, a recent study reported that tumors leverage different pathways to drive EMT, with cancer cells adopting either embryonic- or adult-like programs regulated by distinct transcription factors [47]. Indeed, our RNA-seq findings implicated PRRX2 and TWIST1 as putative transcription factors driving EMT and pro-metastatic phenotypes downstream of CD47 in NSCLC cells, consistent with the embryonic-like EMT trajectory [43].

EMT is a reversible biological process that transforms cells from an epithelial to mesenchymal state which bestows them with motility and invasive capabilities [45]. In physiological conditions, cells undergo EMT to facilitate embryonic development and tissue regeneration but cancer cells co-opt EMT to support stemness, metastasis, and resistance to therapy, all of which drive tumor progression [45,48]. Several studies have reported a correlation between CD47 and EMT, with CD47 expression levels being associated with EMT marker expression, histological features of EMT, and metastasis in tumor tissues from multiple cancer types including colorectal, prostate, and triple negative breast cancers [14,49–52]. Genetic perturbation studies have implicated CD47 in regulating EMT marker expression and phenotypes in ovarian cancer, oral squamous cell carcinoma, and hepatocellular carcinoma models, although the underlying mechanism was not investigated [52–54]. Interestingly, a breast cancer study identified CD47 as a target of the canonical pro-EMT transcription factors, ZEB1 and SNAIL [55]. A later study identified CD47 as a target of ZEB1 in a murine model of Kras-driven lung adenocarcinoma [56]. This group reported that CD47 was expressed on invading lung cancer cells and reprogrammed tumor associated macrophages to protect invading cancer cells from immune destruction [56]. Thus, current evidence indicates CD47 both induces cancer EMT and is itself induced by EMT, suggesting CD47 may be an effector of a feedback mechanism that reinforces EMT programs; however, mechanistic studies are required to investigate this supposition.

Our transcriptomic analyses revealed that CD47 positively regulates molecular programs to promote EMT phenotypes in NSCLC. This discovery presents a significant advance upon previous work in lung and other cancers. Ours is one of few studies to perform RNA-seq on cancer cells with CD47 LOF to understand its influence on cellular programming, and the first to do so in cancer cells engineered with complete CD47 KO. Dong et al. performed RNA-seq on 8505C thyroid cancer cells with CD47 knockdown to investigate correlations between the effects of CD47 and IFT57 on the transcriptome [57]. These authors reported differentially expressed genes but did not perform gene set enrichment analyses (GSEA) to identify pathways influenced by CD47 perturbation. We used EnrichR to perform enrichment analysis of the genes they identified as significantly downregulated upon CD47 knockdown. This identified “Epithelial Mesenchymal Transition” as the second most significant hallmark pathway enriched in their dataset (not shown). In a different study, the same group derived gene expression profiles for triple negative breast cancer cells treated with the CD47 function-blocking agents, SIRPɑ-Fc fusion protein or anti-CD47 monoclonal antibody CC-90002 [58]. They found that EMT signatures were enriched in inhibitor treated cells but the two inhibitors had opposite effects, suggesting that additional studies are needed to resolve the role of CD47 in regulating EMT programs in breast cancer cells. Hu et al. discovered that CD47 overexpression in colon cancer cell lines promoted cell migration and metastasis. Based on gene expression profiling and mechanistic studies, they determined that CD47 promotes metastasis via positive regulation of MAPK signaling and aerobic glycolysis pathways [59]. Thus, multiple independent transcriptomic and functional studies suggest that CD47 regulates EMT and metastasis in different cancer types; however, further research is required to decipher whether CD47 positively or negatively regulates EMT and to deduce its pro-metastatic effectors in specific cancer contexts. Addressing this need, our work yielded novel insights in NSCLC, revealing that CD47 promotes EMT and metastasis through a pathway that drives EMT transcriptional programs.

Our data suggests that CD47 drives EMT-associated phenotypes through an ERK-dependent mechanism. Indeed, we observed that CD47 KO or overexpression caused a decrease or increase in phospho-ERK levels, respectively, in all but one NSCLC cell line (A549). Consistent with this idea, re-expression of ERK in CD47-deficient LLC and H1299 lines enhanced their adhesion and migration abilities, suggesting the CD47-ERK-EMT axis could be generalizable across NSCLC models. CD47 has previously been described as an upstream regulator of several signal transduction proteins, including SRC, FAK, PI3K, BNIP3, PLCγ1, Cdc42, BECN1, and ERK, which regulate a plethora of cellular functions [16]. Its regulation of MAPK signaling in NSCLC is consistent with studies that identified CD47 as a positive regulator of ERK in colorectal cancer, meningioma, and craniopharyngioma models, suggesting CD47-mediated regulation of MAPK signaling could also be a common mechanism in cancer cells [59–61]. In contrast, stimulation of CD47 with the TSP-1 derived peptide, 4N1K, diminished phospho-ERK expression and chemotaxis in smooth muscle cells, further highlighting the complex, context-specific nature of CD47 signaling and its phenotypic effects [62]. Another study reported that ERK signaling induces CD47 expression in melanoma models [63]. This aligns with findings from recent studies that identified the two most prevalent oncogenic drivers in NSCLC, EGFR and KRAS, as positive regulators of CD47 expression [22,64]. In EGFR-driven tumors, CD47 expression was attributed to constitutive activation of the downstream transcription factors c-Myc and NF-κB, as well as CD47 stabilization by SRC-mediated phosphorylation which prevents its degradation [22]. In the context of KRAS, PI3K-STAT3 signaling was found to upregulate CD47 expression by relieving its suppression by miR-34a [64]. These studies and our own suggest a feed-forward mechanism to amplify ERK signaling and CD47 expression may exist in NSCLC cells.

Our discoveries nominate a putative mechanism to explain the involvement of CD47 in NSCLC metastasis involving activation of MAPK signaling and downstream transcription of EMT-related gene expression signatures. Several studies have reported that high CD47 expression is associated with poor prognosis and the incidence of metastasis in NSCLC patients [11–14], indicating the clinical manifestation of the cell and molecular programs we uncovered in preclinical models. Concordantly, we found that CD47 LOF significantly diminished the ability of NSCLC cells to metastasize to the lungs in tail-vein metastasis models in mice. It is important to note that the presence of phagocytes like macrophages in mice could contribute to clearance of CD47 KO cells *in vivo.* As such, the impaired metastasis phenotype and survival benefit we observed for mice injected with CD47 KO cells could be attributable to a monocyte/macrophage-driven engraftment defect rather than reduced pro-metastatic signaling through the cell-intrinsic CD47-ERK-EMT axis we identified. Our observations of lung metastases formed by CD47 KO LLC cells in syngeneic mice and suspected extra-pulmonary metastases in Nude/Nude mice injected with CD47-deficient H1299 cells, suggest that CD47 KO cells are capable of engrafting and that CD47 positively regulates NSCLC metastasis, at least in part, through a cell-intrinsic mechanism. Supporting this interpretation, Hu et al. found that CD47 LOF significantly reduced the metastatic ability of human SW480 colon cancer cells in Nude mice in which macrophages were depleted using clodronate liposomes [59].

Our work highlights the potential for CD47 inhibition to antagonize NSCLC metastasis in patients, which is a major clinical challenge and contributor to tumor progression and mortality. The ability of CD47 to promote oncogenic signaling pathways, drive tumor immune escape, and facilitate EMT-associated drug resistance and cancer cell dissemination, emphasizes the potential for CD47 blockade to be an effective multimodal therapeutic strategy in lung cancer, particularly for EGFR-driven tumors which are enriched for CD47-positivity and refractory to PD-1 blockade [11,12,65]. Numerous agents that disrupt CD47-SIRPɑ interactions have demonstrated promising anti-cancer effects in preclinical tumor models [13,66,67] but their efficacy in clinical trials has been limited by on-target, off-tumor toxicities [68]. This highlights the need for alternative ways to administer CD47 blockade to realize its therapeutic potential, including methods to deliver drugs preferentially to tumor tissues [69] and combination approaches to prime tumors for response to CD47 inhibition [14]. Our finding that CD47 induces phospho-ERK provides a biological rationale for combining CD47 blockade with EGFR/KRAS-targeted therapies to abrogate residual ERK activity and immune escape following EGFR/KRAS inhibition. Accordingly, the combination of EGFR/KRAS- and CD47-targeted therapies was recently shown to exert synergistic effects in preclinical NSCLC models [21].

This study has yielded additional translational insights pertaining to effective strategies for targeting CD47 and its potential utility as a biomarker for informing therapeutic decisions. Given the toxicities associated with current antibody-based modes of delivering CD47 blockade to patients, inhibiting downstream effectors of CD47 signaling could be a more tolerable, clinically tractable approach for abrogating cell autonomous functions of CD47 that drive tumor progression. Our discovery that CD47 LOF sensitizes some NSCLC cells to paclitaxel but not cisplatin suggests microtubule binding agents that disrupt the cytoskeleton could synergize with CD47 blockade, although further research is required to understand which patients could benefit from such combinations. Supporting this idea, a recent study demonstrated enhanced anti-cancer activity when bispecific antibodies blocking P-glycoprotein and CD47 were combined with paclitaxel in multiple drug resistant cancer models [70]. This finding also demonstrates that inhibiting cell-intrinsic CD47 signaling exposes collateral vulnerabilities in NSCLC cells that could be exploited to maximize the anti-tumor efficacy of CD47 blockade, highlighting the need for additional research to characterize them. Moreover, since our work suggests that CD47 promotes NSCLC metastasis *in vivo*, CD47 expression in tumor biopsies could be leveraged to identify early stage patients who would benefit from adjuvant therapy, including CD47 blockade, to antagonize metastatic dissemination and growth of metastatic lesions after the first line of treatment. In agreement with this, several clinical studies have reported positive associations between CD47 expression and metastatic disease in NSCLC, as we previously reviewed [14]. Therefore, continued research is warranted to develop improved strategies for administering CD47 blockade and to investigate its clinical potential for limiting metastasis, as well as the utility of CD47 as a biomarker for informing patient prognosis and therapeutic decisions in the clinic. Notwithstanding, the improved understanding of cell-intrinsic CD47 functions in NSCLC that our work has provided indicates that clinical translation of CD47 inhibitors could greatly benefit patients due to the multi-faceted contributions of CD47 to tumor progression.

We conducted our studies in cell models representing human and murine NSCLC driven by Ras-family oncogenes. Our use of clonally-derived CD47 KO lines ensured complete CD47 LOF was achieved in the cells studied. We observed similar pro-metastatic phenotypes across multiple clonal CD47 KO cell lines as well as cells with polyclonal CD47 knockdown (A549), suggesting our discoveries are not artefacts of clonal cell populations. Importantly, the phenotypes we identified were evident in multiple NSCLC models as well as transcriptomic profiles of NSCLC cell lines and patient tumors, supporting the generalizability and clinical relevance of our findings. However, it is possible that CD47 signaling and the phenotypes it controls differ in lung cancers with distinct proteome expression patterns, genetic drivers, and histologies. This could explain why CD47 KO did not significantly reduce phospho -ERK expression in A549 cells, and explain differences seen for specific phenotypes in our study and other NSCLC studies (eg. Cdc42 as a CD47 effector protein and CD47-mediated regulation of cisplatin sensitivity [13,71]). Characterizing CD47 signaling in cell models enabled us to study cell-autonomous CD47 signaling in a controlled environment; however, future *in vivo* studies are warranted to define how cell-intrinsic CD47 signaling in cancer cells is influenced by extrinsic cues from immune and other stromal cells in the tumor microenvironment.

Although we identified ERK as a putative downstream effector of CD47 signaling in NSCLC cells, the ligands that stimulate CD47 as well as the effectors up- and downstream of ERK remain to be defined. We observed no differences in expression of CD47 signaling effectors reported in other cellular contexts (eg. phospho-SRC, phospho-FAK, Cdc42) between WT and CD47 KO cells. Moreover, the CD47 ligand thrombospondin-1 (TSP-1) did not induce a CD47-dependent induction of phospho-ERK in NSCLC cells (data not shown). Integrin αvβ3 has been reported to interact with and mediate CD47 signaling, but treatment of NSCLC cells with the integrin αvβ3 inhibitor, Cyclo-(-RGDfK), failed to reduce phospho-ERK expression (data not shown). These results suggest neither TSP-1 or integrin αvβ3 are upstream mediators of CD47-ERK signaling in NSCLC cells. Thus, unbiased profiling studies to identify specific receptor-ligand interactions and effectors that control the CD47-ERK-EMT axis are warranted and could nominate therapeutic targets whose inhibition could be leveraged to antagonize this tumor-promoting signaling axis. In summary, we uncovered new insights into the positive regulation of EMT and pro-metastatic phenotypes by CD47 through a putative ERK-dependent transcriptional mechanism in NSCLC. However, further mechanistic studies are needed to define the CD47 interactome and elucidate specific downstream effector proteins that execute the CD47-ERK-EMT signaling pathway in NSCLC cells.

## 5.0 Conclusion

Our work advances knowledge of the cell-intrinsic functions of CD47 in the context of NSCLC. We discovered that CD47 regulates MAPK signaling and EMT at the molecular level to stimulate pro-metastatic tumor-promoting phenotypes, demonstrating that it functions as more than just an immunosuppressive signal. Although a deeper mechanistic understanding of the CD47-ERK-EMT signaling pathway is needed, our findings highlight the multifaceted role that CD47 plays in lung cancer biology and progression, as well as the multimodal potential for CD47-targeted therapy to benefit NSCLC patients. As such, further research to develop improved therapeutic agents for delivering CD47 blockade with better toxicity profiles and anti-tumor activities is warranted.

## 6.0 Data Accessibility

RNA-seq data will be deposited within the Gene Expression Omnibus upon publication.

## 7.0 Author Contributions

AL and KLT were responsible for study conception, experimental design, and manuscript preparation. AL conducted experiments with supervision and assistance from LZ, RH, ZR, AZ, YFW, RS, HN and KLT. AL and KLT performed data analysis. All authors made significant contributions and critically reviewed the manuscript.

## Acknowledgements

The authors are grateful to Drs. Katalin Szaszi, Andras Kapus, Shannon Dunn, and Sunit Das for helpful scientific discussions and experimental suggestions. The authors thank UHT core facility scientists, Drs. Caterina Di Ciano Oliveira, Xiaofeng Lu, Pamela Plant, and Monika Lodyga for technical support and Dr. Boris Hinz for kindly providing reagents. The authors are also grateful to the Broad Institute for making cancer cell line genomics data publicly available in the DepMap Portal (https://depmap.org/portal) and to The Cancer Genome Atlas for sharing genomics data for clinical tumor specimens.

## Abbreviations

NSCLC: non-small cell lung cancer
EMT: epithelial-mesenchymal transition
TSP-1: thrombospondin-1
KO: knockout
RNP: ribonucleoprotein
CDS: coding sequence
IF: Immunofluorescence
SRB: sulforhodamine B
TCA: trichloroacetic acid
DEG: differentially expressed genes
GSEA: geneset enrichment analyses
MAPK: mitogen-activated protein kinase
FDR: false-discovery rate

