## Supplementary Material for "CD47 promotes MAPK and epithelial-to-mesenchymal transition molecular programs to drive pro-metastatic phenotypes in non-small cell lung cancer"

**Supporting Information**

Table S1. PCR primers
Table S2. qPCR primers
Table S3. MSigDB gene sets
Table S4. sgRNA sequence

Supplementary Tables

*Table S1. PCR Primers*

| **Primer** | **Sequence** | **Purpose** |
|---|---|---|
| mmCd47-gDNA-F | AGTTACAGTCTACTGGCTGGTGTG | PCR and sequencing Cd47 CRISPR edits in mouse gDNA |
| mmCd47-gDNA-R | CTTAACTTGGTTTCTGGTTCTTGG | PCR and sequencing Cd47 CRISPR edits in mouse gDNA |
| HsCD47-gDNA-F | CATCCATATTTATACTGCCGTGAC | PCR and sequencing Cd47 CRISPR edits |
| HsCD47-gDNA-R | AAGAAATGGATTCTTCCATACAGC | PCR and sequencing Cd47 CRISPR edits |
| mm.CD47.CDS.Kozak.F | GCCACCATGTGGCCCTTGGCGGCG | Amplify mouse CDS for overexpression system |
| mm.CD47.CDS.R | ACCTATTCCTAGGAGGTTGGATAGTCCTCTGG | Amplify mouse CDS for overexpression system |
| hs.CD47.CDS.Kozak.F | GCCACCATGTGGCCCCTGGTAGCGGCG | Amplify human CDS for overexpression system |
| hs.CD47.CDS.Kozak.R | TCAGTTATTCCTAGGAGGTTGTATAGTCTTCTGATTGG | Amplify human CDS for overexpression system |
| mm.Kozak-Erk2-F | GCCACCATGGCGGCGGCGGCGGCGGCGGG | Amplify mouse CDS for overexpression system |
| mm.Erk2-Stop.R | TTAAGATCTGTATCCTGGCTGGAAT | Amplify mouse CDS for overexpression system |

*Table S2. qPCR Primers*

| Gene | Forward Primer | Reverse Primer |
|---|---|---|
| mm.Hprt1 | TCAGTCAACGGGGGACATAAA | GGGGCTGTACTGCTTAACCAG |
| Hs.HPRT1 | CCTGGCGTCGTGATTAGTGAT | AGACGTTCAGTCCTGTCCATAA |
| mm.Cd47 | CATCCCAGGAGAAAAGCCCG | GGACGTAGCCCAGCACTTGA |
| Hs.CD47 | CGTGTTGTTTCATGGTTTTC | CTCATCCATACCACCGGATCT |

*Table S3. MSigDB gene sets*

| Human Gene Set | MSigDB ID | Collection | Source Species |
|---|---|---|---|
| GOBP_REGULATION_OF_EPITHELIAL_ TO_MESENCHYMAL_TRANSITION | M16783 (GO:0010717) | C5, GO, BP | Homo sapiens |
| GOBP_POSITIVE_REGULATION_OF_ MAPK_CASCADE | M11151 (GO:0043410) | C5, GO, BP | Homo sapiens |
| GOBP_ERK1_AND_ERK2_CASCADE | M16677 (GO:0070371) | C5, GO, BP | Homo sapiens |
| JECHLINGER_EPITHELIAL_TO_ MESENCHYMAL_TRANSITION_UP | M1406 | C2, CGP | Mus musculus |
| KOHN_EMT_MESENCHYMAL | M46417 | C2, CGP | Homo sapiens |

*Table S4. sgRNA sequences*

| Oligo | Guide Sequence |
|---|---|
| mm.sgCd47-F1 | TATAGAGCTGAAAAACCGCA |
| mm.sgCd47-F2 | CCACATTACGGACGATGCAA |
| sgLacZ_F1 | GCCCGAATCTCTATCGTGCGG |
| hs.sgCD47-6F | TAACAGAATTAACCAGAGA |
| hs.sgCD47-7F | AGTGATGCTGTCTCACACAC |
| hs.sgAAVS1-F | GGGGGCCACTAGGGACAGGAT |

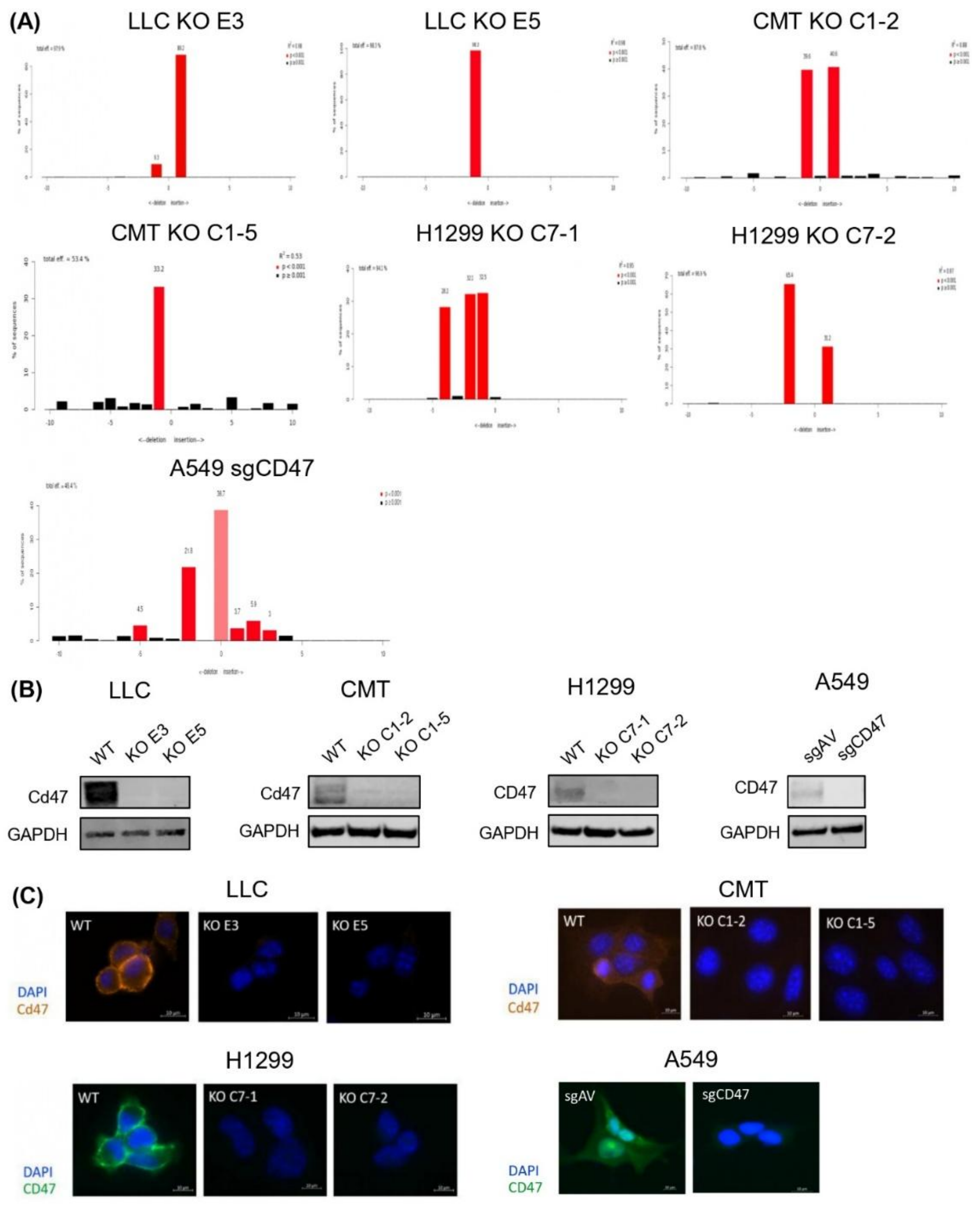


**Figure S1. Confirmation of *CD47* gene editing and loss of protein expression in NSCLC models engineered with CD47 knockout (KO) or knockdown. A)** Detection of CRISPR/Cas9-induced indels generated in clonal LLC, CMT, and H1299 CD47 KO lines. The products of PCR done on genomic DNA spanning the sgRNA target site within the *CD47* gene were subjected to Sanger sequencing. Tracking of Indels by DEcomposition (TIDE) analysis was used to deconvolute the sequences to identify specific indels created in each model. Plots produced by the TIDE deskgen tool are shown for the clonally-derived KO lines (LLC KO E3 and E5, CMT KO C1-2 and C1-5, H1299 KO C7-1 and C7-2) as well as polyclonal A549 cells with CRISPR-mediated CD47 knockdown. Red bars show the fraction of each edited allele within the KO/knockdown lines. The estimated percentage of editing in each model is also indicated. **B,C)** Western blots and immunofluorescence confirming loss of CD47 expression in the 4 isogenic lines.

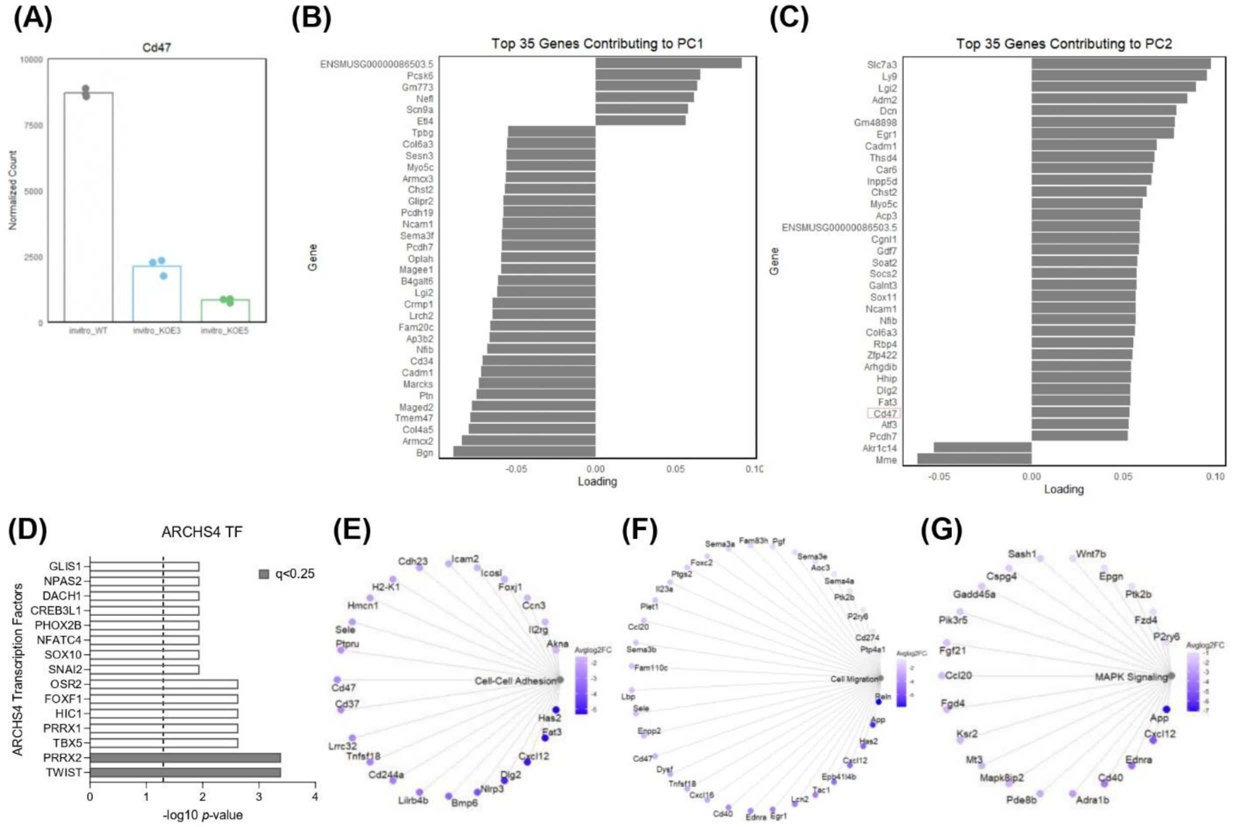


**Figure S2. CD47 expression and genes contributing to the principal components identified based on transcriptomic profiling of LLC WT and CD47 KO cells. A)** Normalized counts for RNA-seq reads mapping to CD47 in WT and CD47 KO LLC cells. **B,C)** The top 35 genes contributing to PC1 (B) and PC2 (C) identified in the principal component analysis. The red box in (C) indicates that Cd47 is one of the PC2 genes. **D)** Top ARCHS4 Transcription Factor gene sets enriched for DEGs downregulated in CD47 KO LLC cells identified using EnrichR. **E-G)** Genes contributing to the Cell-Cell Adhesion, Cell Migration, and MAPK Signaling processes enriched for DEGs that are downregulated in CD47 KO cells (Fig.2F-G). The colour scale indicates the average $\log_2$ fold change in expression across both CD47 KO clones relative to WT cells.

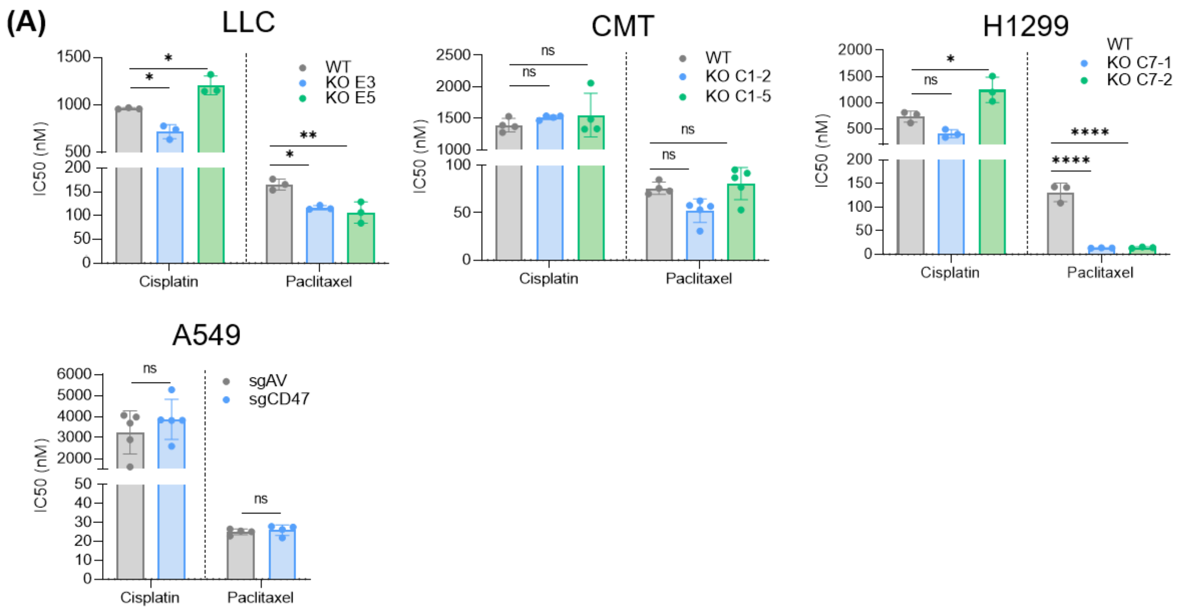


**Figure S3. Effect of CD47 loss of function on chemotherapy response in NSCLC lines. A)** Dose response assays were conducted in LLC, CMT, H1299, and A549 cells in 96-well plates. WT and KO cells were plated and treated the next day with 9 serial dilutions of cisplatin (50µM, 3-fold dilutions) or paclitaxel (500nM, 2-fold). After 5 days, surviving cells were fixed with sulforhodamine B (SRB). SRB was solubilized with Tris and absorbance at 570nm was read on a microplate spectrophotometer as a readout of cell viability. Viability for each drug dose was normalized to vehicle treated controls. Drug dose response curves were plotted and analyzed in GraphPad to calculate $IC_{50}$ values. $IC_{50}$ values were compared between WT and CD47 KO cells using a two-tailed student's t-test. Each data point indicates one experimental replicate. Error bars indicate mean ± SD.

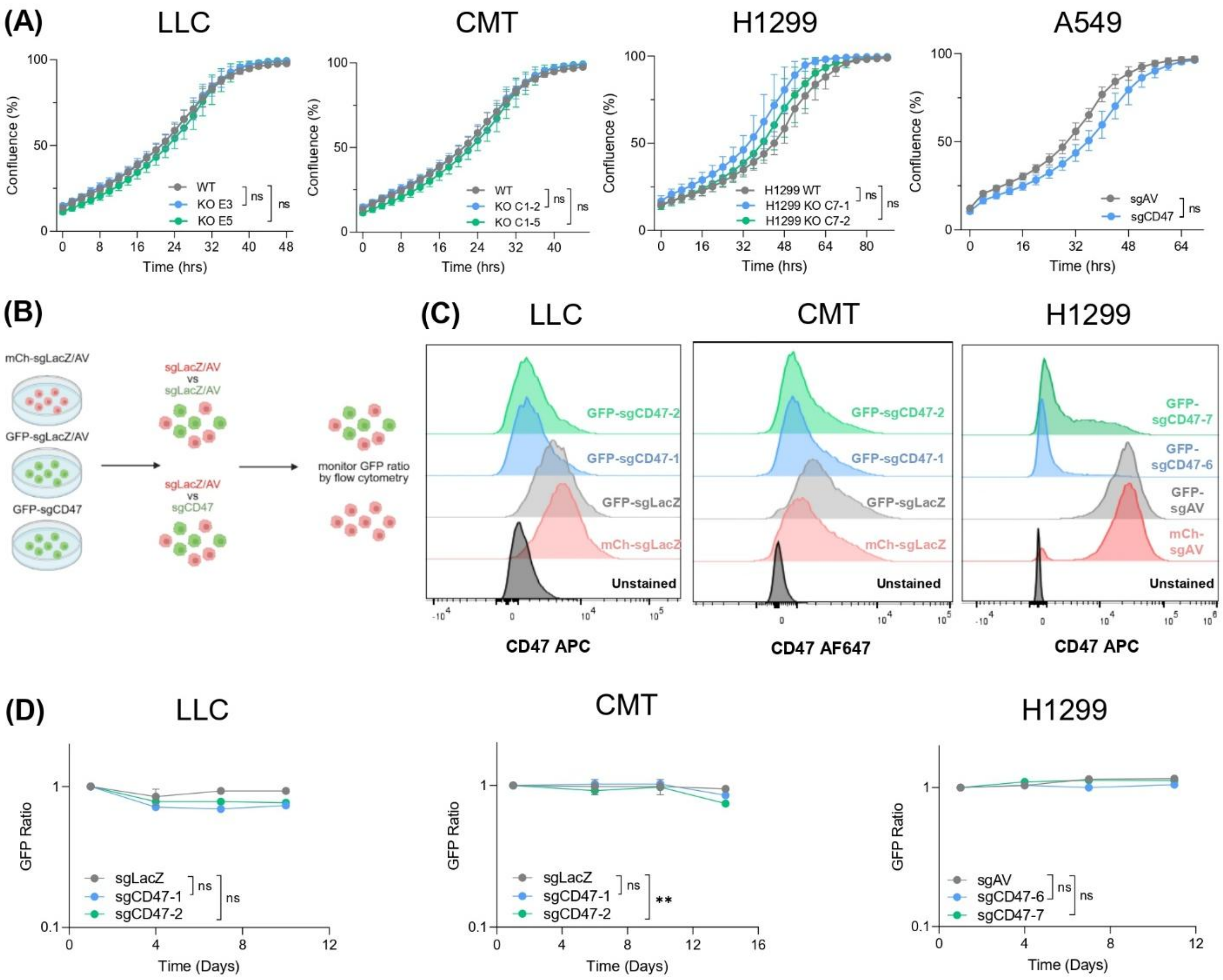


**Figure S4. CD47 KO does not affect lung cancer cell proliferation or fitness. A)** Imaging-based proliferation assay monitoring cell confluence over 48-92 hours. Cells were plated sparsely in 24-well plates and imaged every 2 hours. Percent confluence indicates the fraction of well area covered by cells. Comparisons of confluencies between WT and CD47 KO cells were done using two-tailed student's t-test with multiple testing correction across all time points for each cell line. T-tests between WT and KO cells were non-significant at every time point. **B)** Schematic representation of the multi-colour competition assays. Assay details are described in the methods. Briefly, Cas9+ LLC, CMT and H1299 cells were transduced with mCherry (mCH) or GFP lentivectors to express control sgRNA targeting LacZ (for mouse) or AAVS1 (AV, for human), or sgRNA targeting CD47 (CD47-1, CD47-2). GFP+ sgCD47 cells were mixed 1:1 with GFP+ or mCH+ sgAAVS1/sgLacZ control cells and the ratio of GFP:mCH cells was monitored over several passages with flow cytometry. **C)** Loss of CD47 surface expression in sgCD47 compared to control cell populations confirmed by flow cytometry. **D)** Results of assays competing cells with WT (sgLacZ, sgAV) *CD47* versus cells with genetically disrupted *CD47* (sgCD47) indicating no appreciable effect of CD47 perturbation on NSCLC cell fitness. GFP+ ratios between the different cell mixtures were compared at the final passage using a student's t-test. Data are representative of three independent experiments for each model.

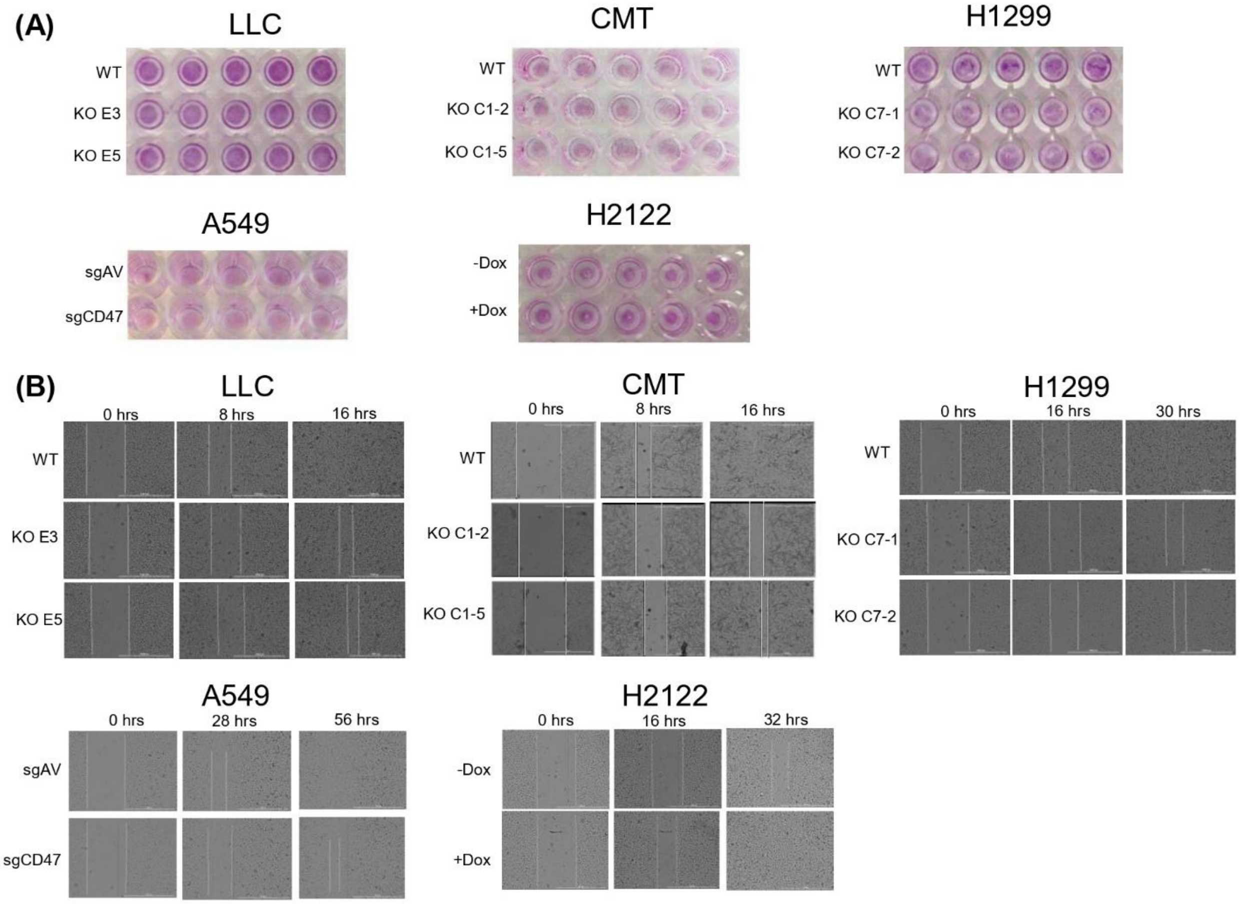


**Figure S5. Representative images for cell adhesion and migration assays in NSCLC cells.** Representative cell adhesion (**A**) and migration (**B**) results for experiments presented in Fig.4. Pink sulforhodamine B staining for adhesion assays was quantified using absorbance at 570nm on a spectrophotometer. Vertical lines in the migration assays indicate wound boundaries which were used to quantify wound width at the time points indicated.

(A)

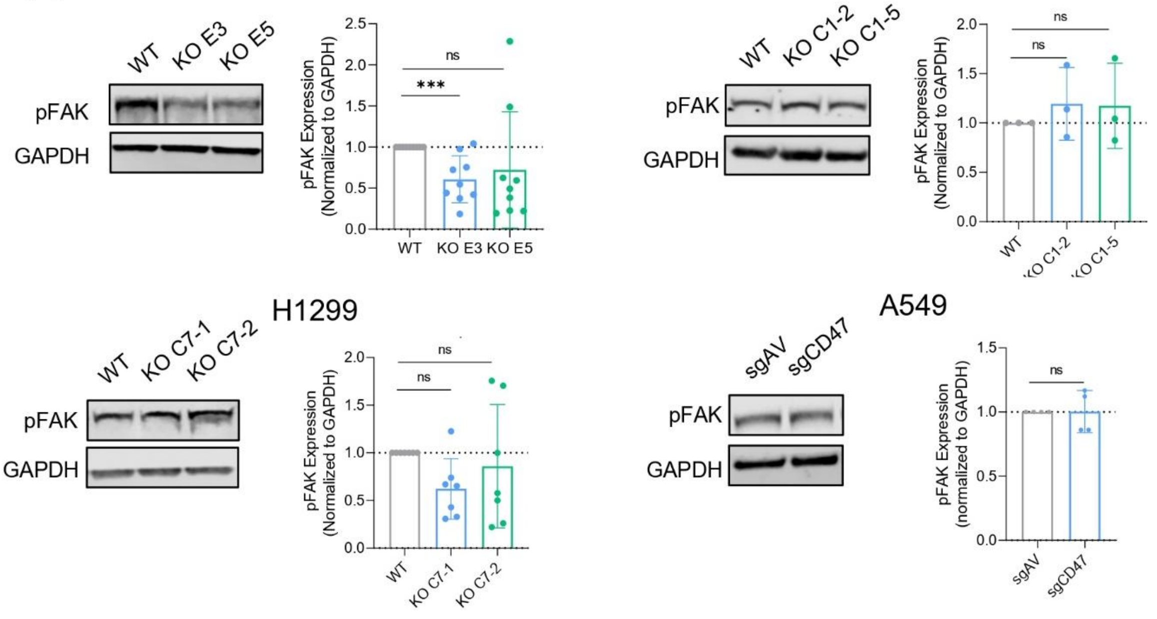
LLC
WT
KO E3
KO E5
pFAK
GAPDH
pFAK Expression
(Normalized to GAPDH)
***
ns
CMT
KO C1-2
KO C1-5
H1299
KO C7-1
KO C7-2
A549
sgAV
sgCD47
(normalized to GAPDH)

(B)

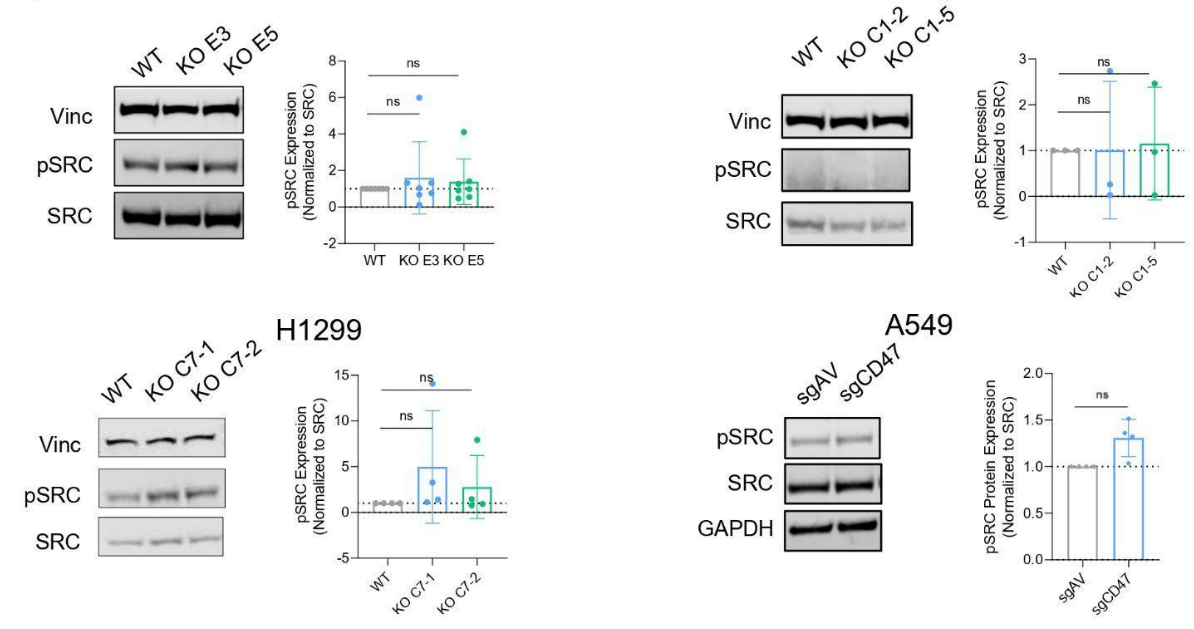
LLC
WT
KO E3
KO E5
Vinc
pSRC
SRC
pSRC Expression
(Normalized to SRC)
ns
CMT
KO C1-2
KO C1-5
H1299
KO C7-1
KO C7-2
A549
sgAV
sgCD47
GAPDH
pSRC Protein Expression
(Normalized to SRC)

(C)

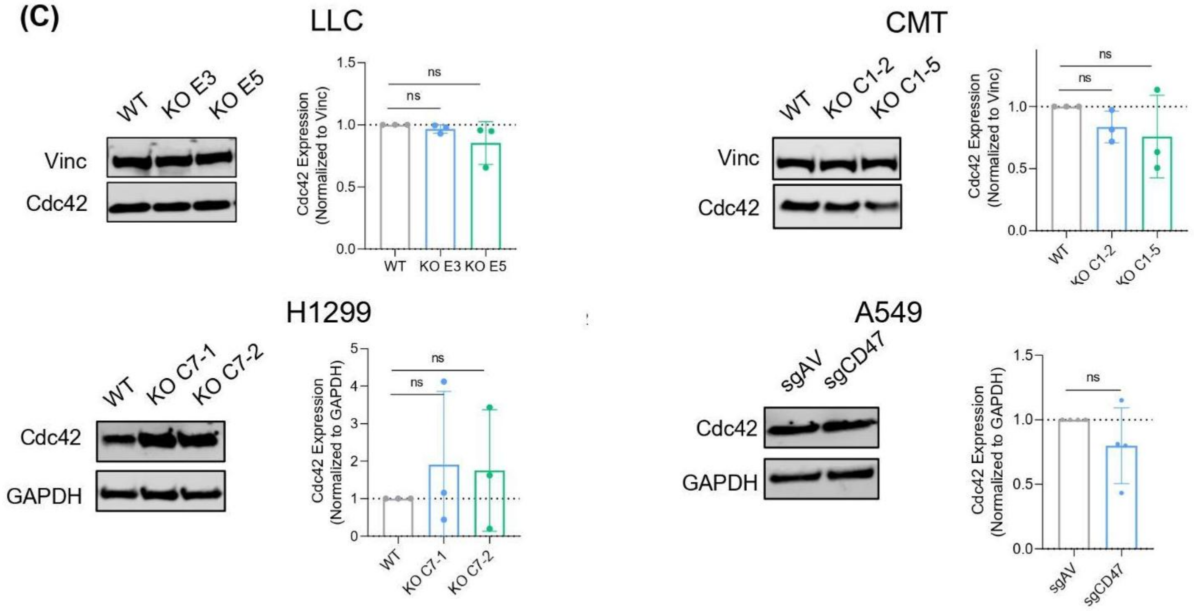


**Figure S6. Expression of reported CD47 effectors, FAK, SRC, and Cdc42 in WT and CD47 KO/knockdown cells.** Representative western blots results for phospho-FAK (A), phospho- and total SRC (B), and Cdc42 (C) in isogenic LLC, CMT, H1299 and A549 cells. Note, the GAPDH control for H1299 cells in (A) is the same as that in Fig.5C. For quantitation, pSRC was normalized to total SRC, and phospho-FAK and Cdc42 were normalized to Vinculin (Vinc) or GAPDH. Each data point indicates one biological replicate. A one sample t-test was done on fold change data to detect differences in protein expression between cells with CD47 perturbation and WT controls. No reproducible changes in potential CD47 effector proteins were observed in KO cells compared to WT controls across the models suggesting CD47 signals through other effector proteins. Error bars indicate mean ± SD.

Figure S7

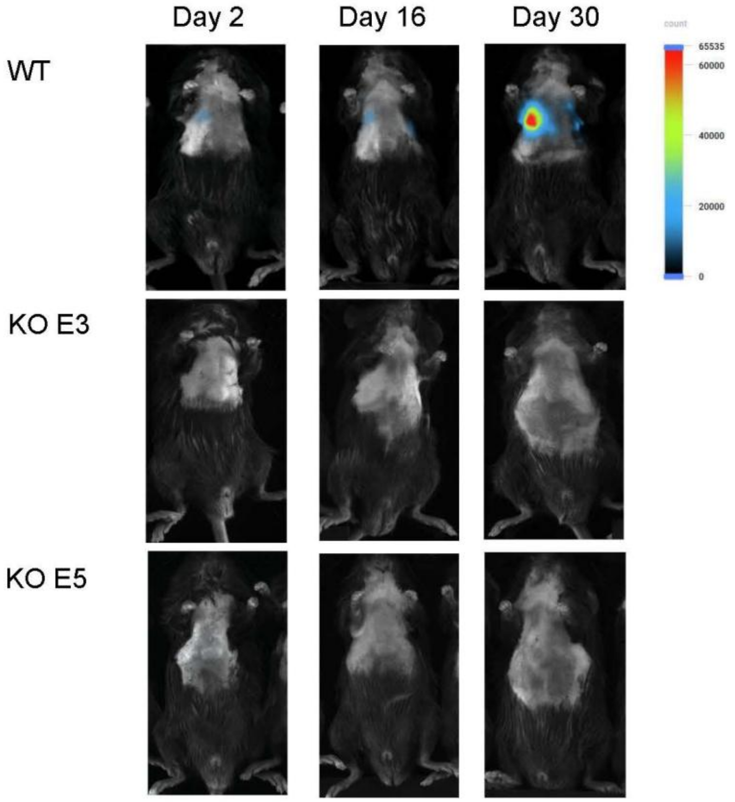


**Figure S7. Representative bioluminescence images of mice injected with LLC-Luc cells.** C57BL/6 mice were intravenously injected (tail vein) with LLC-Luc cells. From Day 2 onwards, mice were imaged with BLI every 2-3 days to monitor for tumor formation.
